# Effectiveness of protected areas in conserving tropical forest birds

**DOI:** 10.1101/2020.01.21.912345

**Authors:** Victor Cazalis, Karine Princé, Jean-Baptiste Mihoub, Joseph Kelly, Stuart H.M. Butchart, Ana S.L. Rodrigues

## Abstract

Protected areas are the cornerstones of global biodiversity conservation efforts^1,2^, but to fulfil this role they must be effective at conserving the ecosystems and species that occur within their boundaries. This is particularly imperative in tropical forest hotspots, regions that concentrate a major fraction of the world’s biodiversity while also being under intense human pressure^3–5^. But these areas strongly lack adequate monitoring datasets enabling to contrast biodiversity in protected areas with comparable unprotected sites^6,7^. Here we take advantage of the world’s largest citizen science biodiversity dataset – eBird^8^ – to quantify the extent to which protected areas in eight tropical forest biodiversity hotspots are effective at retaining bird diversity, and to understand the underlying mechanisms. We found generally positive effects of protection on the diversity of bird species that are forest-dependent, endemic to the hotspots, or threatened or Near Threatened, but not on overall bird species richness. Furthermore, we show that in most of the hotspots examined this is driven by protected areas preventing both forest loss and degradation. Our results support calls for increasing the extent and strengthening the management efforts within protected areas to reduce global biodiversity loss^9–11^.

---

Hopes for halting and reversing the ongoing global biodiversity crisis are largely pinned on protected areas^1,12^. Defined as geographical spaces recognised, dedicated and managed to achieve the long term conservation of nature^1^, they are expected to buffer ecosystems and species populations against some of the most destructive impacts of human activities, particularly those resulting in habitat loss or degradation, or the overexploitation of wildlife. Already covering nearly 15% of the global land surface and 7.8% of the oceans^1^, signatories to the Convention on Biological Diversity have committed through Aichi Target 11 to expanding protected area coverage to respectively 17% and 10% by 2020^13^, and there are calls to go much further^14^. However protected areas can only fulfil their intended role if they are effective.

## Challenges to assessing protected area effectiveness

Protected area effectiveness can be assessed through multiple, complementary approaches, for instance, by evaluating whether they cover the diversity of species and ecosystems and the most important sites, or by assessing their management adequacy in terms of staff or resources^1,15^. Here, we focus on effectiveness in terms of biodiversity outcomes: the extent to which the establishment of protected areas makes a difference to the trends and thus ultimately to the condition of the species and ecosystems within their boundaries.

Evaluating outcomes is not straightforward, because it requires contrasting current state with a counterfactual, i.e., an alternative scenario of what would have happened if the protected area had not existed^15^. Simply contrasting any protected and unprotected sites would not be an adequate counterfactual analysis, because it would conflate implementation effects (the difference protected areas have made) with location biases (differences between protected and unprotected sites prior to protected areas implementation)^15,16^. Such location biases are inevitable because protected areas tend to be placed in regions of little economic interest (i.e. greater remoteness, higher altitudes, and lower agricultural suitability^16,17^), which are less likely to have suffered from human pressure both prior and after protection. These differences can be statistically controlled for in counterfactual analyses of protected area effectiveness^6,7,18^, however this requires large datasets on the spatial distribution of the biodiversity features of interest across many protected and unprotected sites.

Nowhere are effective protected areas more essential than in tropical regions, which host a major share of the world’s biodiversity^3^ and are facing rapid habitat loss^3^ and degradation^4,19^, both considered main threats to biodiversity^19–21^.Yet, evaluating protected area effectiveness in these regions is particularly challenging, given that the detailed biodiversity datasets required for counterfactual analyses are typically unavailable^22^. Among the few analyses investigating biodiversity outcomes of tropical protected areas, most focused on protected areas effects on habitats, finding that they mitigate both forest loss and forest degradation^18,23–25^. While such analyses can utilise remote sensing data, investigating effectiveness in terms of species outcomes essentially requires data collected *in situ*. Two global meta-analyses reviewed local-scale studies that had contrasted protected versus unprotected sites in terms of species diversity^7,26^. Both uncovered positive effects at the global scale, but – worryingly – weaker or mixed results within tropical regions, contrasting with reported positive effects of protected areas at reducing forest loss and degradation.

## Assessing effectiveness in tropical forest hotspots

In this study, we investigate the effectiveness of protected areas in eight tropical forest biodiversity hotspots, using a counterfactual analysis that controls for location biases to quantify outcomes in terms of bird species diversity. For this purpose, we take advantage of an exceptional recent dataset on the fine-scale occurrence of bird species, built through eBird, the world’s largest citizen science programme^8^. Even though eBird records are not collected through a standardised protocol, the sheer size of the database allows us to control for confounding effects (observer experience, sampling effort, seasonality) that can affect the length and composition of recorded bird lists^27^.

We focus on biodiversity hotspots, i.e., biogeographic regions with high levels of species endemism that have lost most of their original habitat^5^. These are the epicentres of the ongoing biodiversity crisis^3,28^, and thus regions where effective conservation efforts are the most urgent. We analysed eight tropical forest hotspots with good coverage by eBird observations^29^ (i.e. with >1,000 sampling events per hotspot after data filtering): four in the Americas (Atlantic Forest, Tropical Andes, Tumbes-Chocó-Magdalena, Mesoamerica), one in Africa (Eastern Afromontane), and three in Asia (Western Ghats and Sri Lanka, Indo-Burma, Sundaland). We focused on the area within each hotspot included in the “Tropical and subtropical moist broadleaf forests” biome^30^ (Fig. 1, Extended Fig. 1), assumed to have been originally forested (Supplementary Methods 4D).

**Fig. 1:**
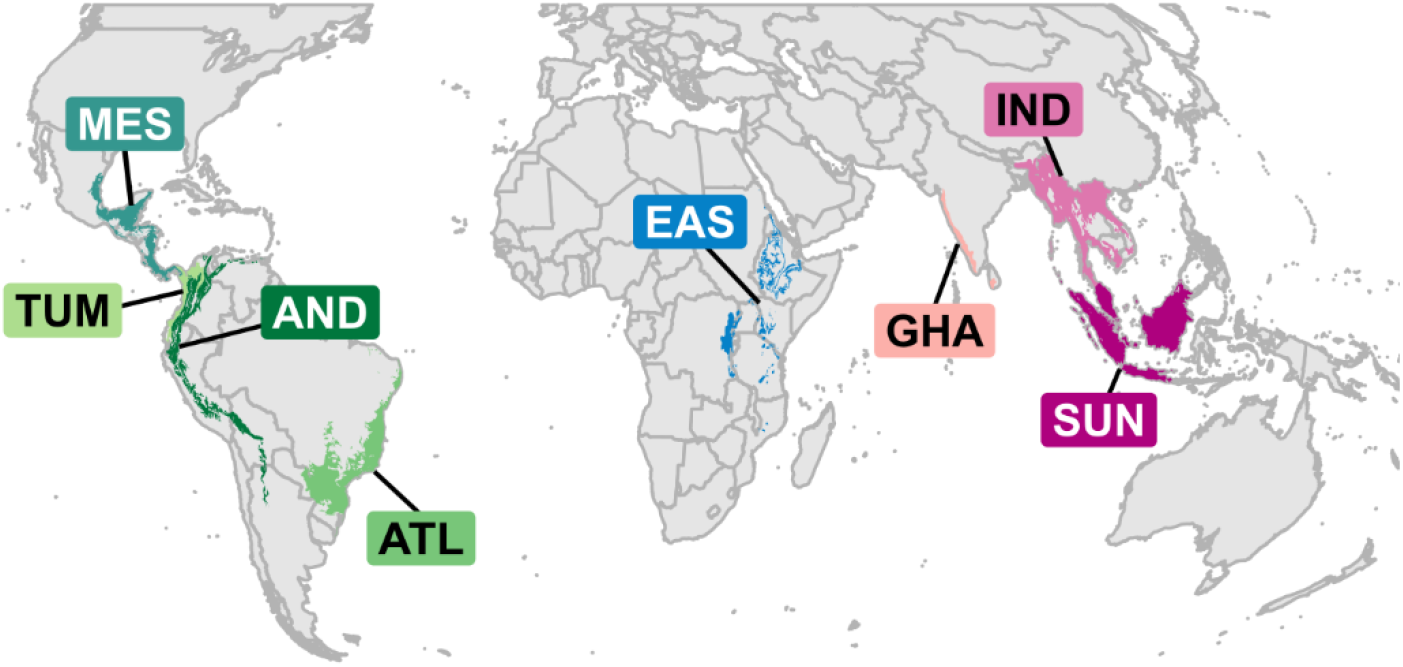
Regions covered by the present study (i.e. intersection between eight biodiversity hotspots and the “tropical and subtropical moist broadleaf forests” biome). Acronyms and colours as in Fig. 3 and Fig. 4: ATL (Atlantic Forest, N=6,760 checklists), AND (Tropical Andes, N=17,758), TUM (Tumbes-Chocó-Magdalena, N=1,188), MES (Mesoamerica, N=32,784), EAS (Eastern Afromontane, N=1,097), GHA (Western Ghats and Sri Lanka, N=2,646), IND (Indo-Burma, N=2,996), SUN (Sundaland, N=1,548). More details in Extended Fig. 1.

For each hotspot, we apply a set of three distinct but interrelated statistical analyses to investigate the effectiveness of protected areas at retaining bird diversity and to shed light on the underlying mechanisms (Fig. 2). First (analysis I), we test whether protected areas differ from unprotected sites in terms of their bird species diversity, after controlling for location biases (altitude, remoteness, and agricultural suitability of sites) (Fig. 3). We quantify bird species diversity using four indices, one being overall species richness and the others richness in three (partially overlapping) types of species of conservation concern, namely: specialists (here, forest-dependent species); species with narrow ranges (i.e., endemic to the hotspot); and species classified as threatened (Critically Endangered, Endangered, or Vulnerable) or Near Threatened in the IUCN Red List^31^. We consider two potential mechanisms through which protected areas can potentially affect bird diversity: by retaining forest presence (i.e. mitigating forest loss); and by maintaining forest quality (i.e., mitigating forest degradation). We test these mechanisms in two complementary analyses (Fig. 2). One (analysis II) tests the effects of protected areas on forest presence (IIa) or on each of three measures of forest quality (IIb): canopy height; forest contiguity (i.e., the opposite of fragmentation); and wilderness (i.e., the opposite of the human footprint index^32^). The other (analysis III) tests the effects of either forest presence (IIIa) or forest quality (IIIb) on each of the four above-mentioned indices of bird diversity. In the latter analysis (IIIb), we also considered the residual effects of protected areas beyond the three variables of forest quality, to account for a possible effect of protection on other forms of habitat degradation (e.g., hunting, understorey thinning, invasive species) that we could not measure directly (Fig. 4).

**Fig. 2:**
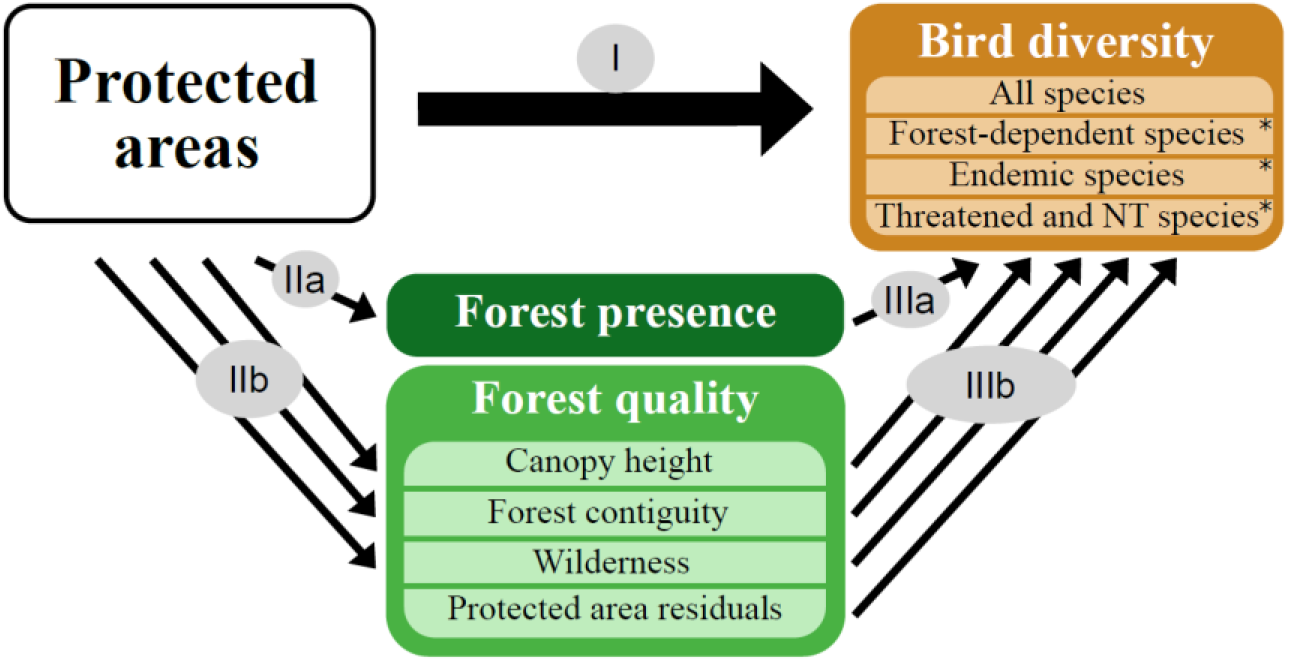
Framework of the analyses performed to investigate the effectiveness of protected areas at retaining bird diversity. Analysis I: effect of protected areas on bird diversity measured through four indices of bird species richness (all species, forest-dependent species, endemic species, threatened and Near Threatened species). The asterisk indicates species of conservation concern. Analysis II: effects of protected areas on forest presence (IIa) and on three measures of forest quality (canopy height, forest contiguity, and wilderness; IIb). Analysis III: effects of forest presence (IIIa), and of each of the three measures of forest quality and of the residual effect of protected areas (IIIb) on bird diversity.

**Fig. 3:**
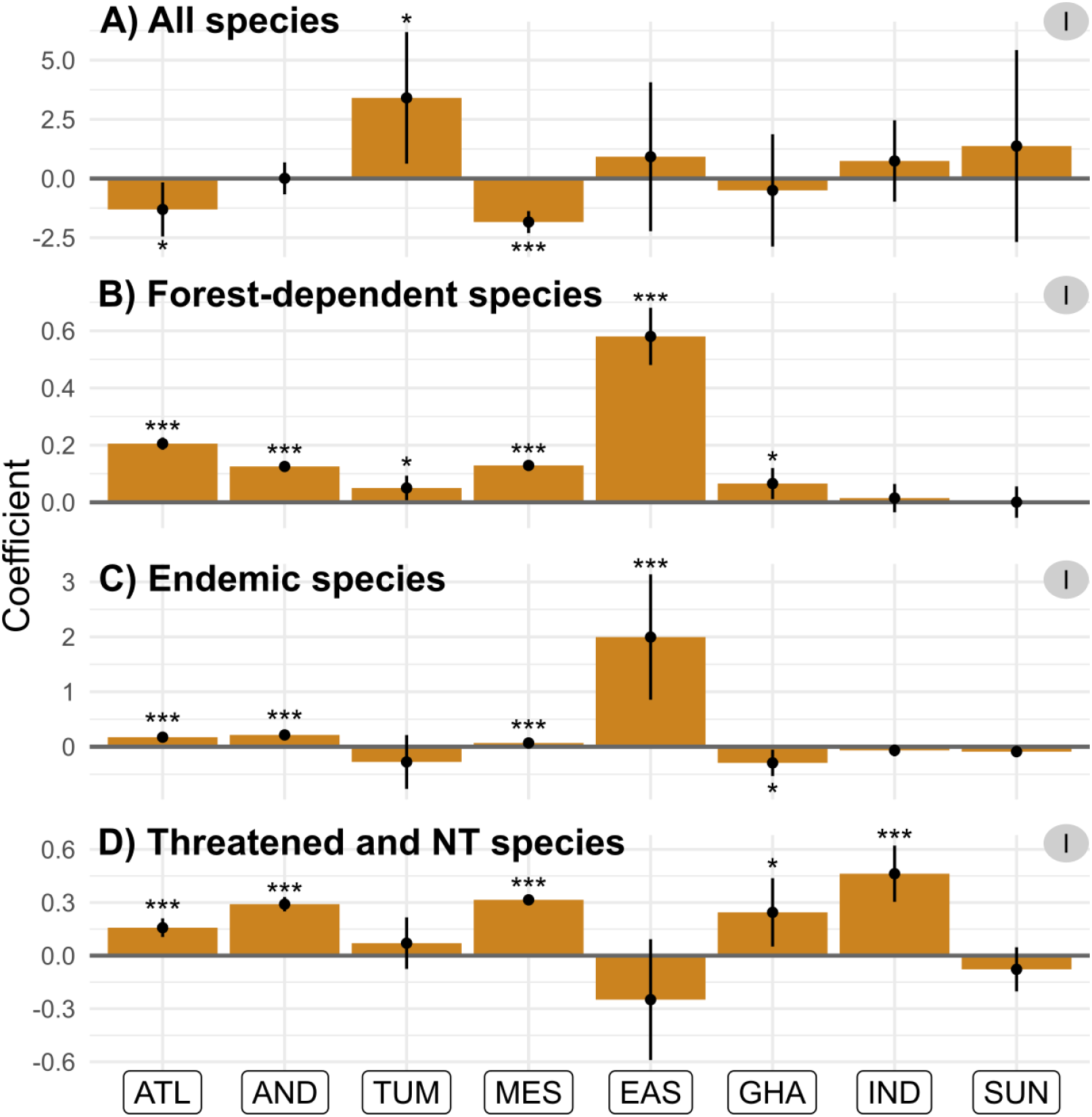
Effect of protected areas on bird diversity per hotspot, for four bird diversity indices (analysis I). A) overall species richness; B) forest-dependent species richness; C) endemic species richness; D) richness in threatened and Near Threatened species. Coefficients correspond to the estimates of GAM models; significance is given by the P-value (***<0.001<**<0.10<*<0.05) and the 95% confidence interval (vertical lines). Hotspots: ATL (Atlantic Forest), AND (Tropical Andes), TUM (Tumbes-Chocó-Magdalena), MES (Mesoamerica), EAS (Eastern Afromontane), GHA (Western Ghats and Sri Lanka), IND (Indo-Burma), SUN (Sundaland).

**Fig. 4:**
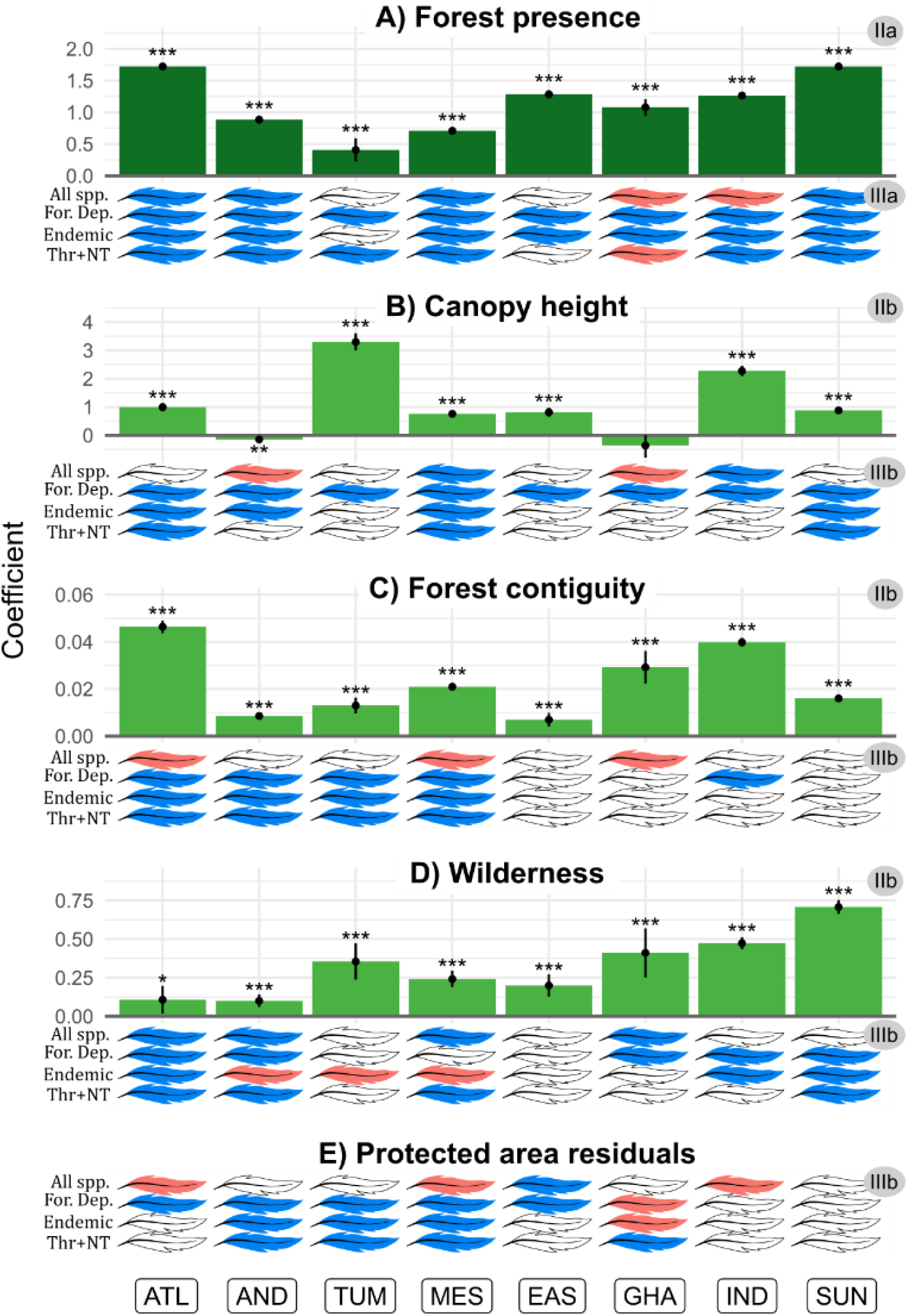
Effects of protected areas on habitat, and effects of habitat on bird diversity, per hotspot. Bars: effects of protection on forest presence (A; analysis IIa) or on forest quality (B, canopy height; C, forest contiguity; D, wilderness; E, protected area residuals; analysis IIb); coefficients correspond to the estimates of GAM models; significance given by P-value (***<0.001<**<0.10<*<0.05), and 95% confidence interval (vertical lines). Feathers: colour represents the effect sign (blue: positive; red: negative; white: non-significant) of each habitat variable on each of the bird diversity variables (All spp., overall species richness; For.Dep., richness in forest-dependent species; Endemic, richness in endemic species; Thr+NT, richness in threatened and Near Threatened species). Hotspots: ATL (Atlantic Forest), AND (Tropical Andes), TUM (Tumbes-Chocó-Magdalena), MES (Mesoamerica), EAS (Eastern Afromontane), GHA (Western Ghats and Sri Lanka), IND (Indo-Burma), SUN (Sundaland).

By controlling for altitude, remoteness and agricultural suitability of sites, we assume that observed differences between protected and unprotected sites (in analyses I and II) are not a consequence of pre-existing differences when protected areas were established, and instead reflect the effects of protected area implementation on pressures affecting trends in (and thus over time the state of) bird diversity, forest cover or forest quality.

## Protected areas retain species of concern, not all species

We found no consistent evidence of an effect of protected areas on overall richness in bird species (analysis I). Indeed, we obtained non-significant results for five out of the eight hotspots tested, significant negative effects for two, and a significant positive effect for a single one (Fig. 3A). Given that species richness is an intuitive and widely used measure of biodiversity^33^, these results may appear worrying, by suggesting that protected areas do not prevent local biodiversity loss. In fact, they agree with a wealth of previous evidence that overall species richness is not a suitable indicator of local biodiversity impact, as species that go locally extinct due to ecosystem alteration are often replaced by others – often of lower conservation concern – with no impact on overall species richness^33,34^. Accordingly, we also found that neither forest presence (Fig. 4A; Extended Fig. 2A; analysis IIIa) nor forest quality (Fig. 4B-D; Extended Fig. 2B-D; analysis IIIb) had a consistent positive effect on overall species richness, indicating that this diversity measure is rather insensitive to habitat loss and degradation, at least at the temporal and spatial scales considered in this study. Species richness is also known to often increase at intermediate levels of disturbance^35^ perhaps explaining the few observed negative effects of protection (Fig. 3A), of forest presence (Fig. 4A), or forest quality (Fig. 4B-E) on overall species richness.

Whereas we found no effect of protected areas on overall species richness, our results indicate that protected areas are effective at retaining the three types of species of conservation concern analysed: forest-dependent (i.e. specialists), endemics to each hotspot (i.e. narrow-ranged), and threatened or Near Threatened (i.e., at greater risk of extinction). Indeed, for each of these three groups we found significant positive effects of protected areas across hotspots (Fig. 3B-D; analysis IIIb), particularly for forest-dependent species (in 6 out of 8 hotspots; with protected sites on average 17.8% richer in forest-dependent species than comparable unprotected sites; Fig. 3B), but also for endemic species (4/8; 77.6%; Fig. 3C) and threatened and Near Threatened species (5/8; 19.0%; Fig. 3D; Extended Table 1).

Our results corroborate studies in temperate regions that found that protected areas do not protect all species and thus do not always affect species richness^6,36^. However, they contrast with what was known in the tropical regions based on two previous global-scale studies of protected area effectiveness, based on the meta-analysis of local-scale studies contrasting protected *versus* unprotected sites. One of these studies found higher species richness and abundances within protected areas in Africa and Asia but not in South America^26^; the other found higher overall richness within protected areas, but no significant effects on species richness in rare and endemic species, including in the tropics^7^. Nonetheless, the present study provides a stronger test of protected area effectiveness in tropical forests by focusing specifically on these biomes, using more comparable data (as emerging from a single, coherent dataset), better controlling for confounding variables, and by exploring the underlying mechanisms of habitat loss and degradation.

## Protected areas retain diversity in species of concern by mitigating forest loss

Our results suggest that protected areas effectiveness at retaining species of concern is mainly driven by their effectiveness at mitigating forest loss. First, we found significant positive effects of protection on forest presence across all hotspots analysed, with a protected site having on average 17.8% higher probability of being forested than a non-protected counterfactual (Fig. 4A; analysis IIa; Extended Table 1). These results confirm and extend previous works showing positive effects of protected areas at reducing rates of tropical deforestation^18,23,37^. Second, we found that forested sites have higher diversity in forest-dependent bird species than comparable non-forested sites (across 8/8 hotspots; on average 74.9% more species), as well as in endemic species (7/8; 250.0%) and in threatened and Near Threatened species (6/8; 122.1%; analysis IIIa; Extended Table 1; Extended Fig. 2A), in accordance with the well-known devastating impact of deforestation on biodiversity^3,19,21^. Particularly in line with our results, Rutt et al.^38^, have highlighted the replacement of forest-dependent bird species by generalist species following an experimental deforestation in the Amazon.

## Protected areas retain diversity in species of concern by mitigating forest degradation

Our results further indicate that the added value of protected areas towards the conservation of species of concern also comes from their effect at mitigating forest degradation. Firstly, we found a general positive effect of protection on forest quality (analysis IIb), as measured through each of three measures: canopy height (6/8 hotspots; on average 4.8% higher in protected than in counterfactual forested non-protected sites; Fig. 4B), forest contiguity (8/8; 2.6% higher; Fig. 4C), and wilderness (8/8; 5.7% higher; Fig. 4D; Extended Table 1). The last is the reciprocal result of two recent studies showing lower levels of human pressures within protected areas when compared with appropriate counterfactuals in tropical forests^24,25^, whereas the first two results are new contributions to the literature on protected area effectiveness in general, and in tropical regions in particular.

Secondly, our results suggest that each of these three variables of habitat quality enhances diversity in species of concern. Indeed, we show a positive effect of canopy height (in 8/8 hotspots for richness in forest-dependent species; 4/8 for endemic species; 3/8 for threatened and Near Threatened species; Fig. 4B; analysis IIIb; Extended Fig. 2B), of forest contiguity (in 5/8 hotspots for forest-dependent; 4/8 for endemic and for threatened and Near Threatened species; Fig. 4C; analysis IIIb; Extended Fig. 2C) and of wilderness (in 5/8 for forest-dependent species; 3/8 for endemics [but also 3/8 negative]; 4/8 for threatened and Near Threatened species; Fig. 4D; analysis IIIb; Extended Fig. 2D). Finally, even after controlling for canopy height, contiguity, and wilderness, we found that among forested protected areas there are generally positive residual effects of protection itself on forest-dependent species (positive in 5/8 hotspots; but 1/8 negative), on endemics (3/8 positive; but 1/8 negative) and on threatened and Near Threatened species (4/8 positive) (Fig. 4E; Extended Fig. 2E). This indicates that the positive effect of protection in mitigating forest degradation goes beyond the three habitat quality variables we have considered, perhaps reflecting reductions in other types of pressures such as hunting, selective logging, or invasive species^39,40^.

## Stronger evidence of effectiveness for South American protected areas

We found substantial variability across hotspots in the effects of protected areas on the diversity of species of concern. Indeed, the most consistent picture emerges for three of the American hotspots – Atlantic Forest (ATL), Tropical Andes (AND) and Mesoamerica (MES) – for which we found consistently significant positive effects of protection on the three groups of species of concern (Fig. 3), with both forest presence (Fig. 4A) and forest quality (Fig. 4B-E) playing seemingly important roles. Results were more mixed for the other hotspots. We found significant effects of protection on the diversity of forest-dependent species for the Tumbes-Chocó-Magdalena hotspot (TUM), of forest-dependent and endemic species for the Eastern Afromontane hotspot (EAS), of forest-dependent species and threatened and Near Threatened – as well as a negative effect on endemic species – for the Western Ghats and Sri Lanka (GHA), of species of concern for Indo-Burma (IND), and no significant effects on species of conservation concern for Sundaland (SUN). This may reflect variation in the effectiveness of protected area implementation across the world, or simply differences in statistical power. Indeed, the three American hotspots with the strongest signal of effectiveness are those with the most data (6,760-32,784 checklists, contrasted with 1,097-2,996 for the other hotspots; Extended Fig. 1, Extended Fig. 3–4; Supplementary Discussion).

## Conclusions

We provide evidence for the effectiveness of protected areas as biodiversity conservation tools across eight global biodiversity hotspots, covering some of the Planet’s most diverse and threatened terrestrial ecosystems^5^. Through a counterfactual analysis that controls for location biases in the establishment of protected areas, we aimed to isolate as much as possible the effects of implementation itself, i.e., the added value of protection. We found that this value does not lie in preventing declines in overall local species richness, but in avoiding the replacement of species that are most in need of conservation efforts: the forest specialists that are most at risk from forest loss or degradation; the endemic species that make each hotspot globally irreplaceable; and threatened or Near Threatened that are at higher risk of global extinction.

Our results contribute to the body of evidence supporting the effectiveness of protected areas at avoiding forest loss^18,23^, now specifically within the context of tropical forests within biodiversity hotspots. Furthermore, they indicate that this is the main mechanism through which protection has a positive effect on retaining bird species of concern. In addition, we provide evidence that it is not the only mechanism, with protection also having a significant effect on bird diversity by mitigating forest degradation, as measured through canopy height, fragmentation and wilderness levels. Finally, we found evidence for a residual effect of protection (once controlling for the effects on forest presence and quality) that may reflect management measures of other pressures such as hunting, small-scale logging, or invasive species.

In this study, we found that protected areas are effective in the sense that they perform better than comparable unprotected sites. We have, however, not demonstrated that they are sufficiently effective to halt habitat loss and degradation (which previous studies found to be ongoing and sometimes increasing within protected areas^23–25^) nor that they halt population declines (which are still ongoing within many protected areas^41–43^). Furthermore, our analysis does not address whether protected areas are sufficient in terms of their extent of coverage or their representativeness (while previous studies attest that they are not^2,44^). Nonetheless, our results indicate that protected areas are already making a measurable difference in terms of biodiversity conservation in several regions of the world where the conservation stakes are the highest. In this year of 2020 when Aichi Targets are due to be reached^13^ yet some governments are announcing protected areas degazettement and downsizing^45^, our results support the key role of protected areas as global biodiversity conservation tools. We thus join calls for the strategic expansion of the global protected areas estate and increased investments to ensure that they are effectively managed^9–11^.

## Supporting information

Supplementary discussion

Species taxonomy spreadsheet

## Methods

### Study areas: biodiversity hotspots

We focused on eight biodiversity hotspots^5^: those with at least 25% of their extent within the “Tropical and subtropical moist broadleaf forests” biome^30^ and for which we obtained at least 1,000 checklists from eBird (after applying the data selection procedure described below): Atlantic Forest, Tropical Andes, Tumbes-Chocó-Magdalena, and Mesoamerica (Americas); Eastern Afromontane (Africa); Western Ghats and Sri Lanka, Indo-Burma and Sundaland (Asia). Within each hotspot, we analysed only areas overlapping the “Tropical and subtropical moist broadleaf forests” biome^30^ (Fig. 1, Extended Fig. 1; Extended Fig. 3–4), assumed to have been originally forested (see Supplementary Methods 4D).

### Data selection: eBird checklists

We obtained bird sightings from the eBird citizen science database^8^. The reporting system is based on checklists, whereby the observer provides: list of birds detected; GPS location; sampling effort (whether or not all detected species are reported; sampling duration; sampling protocol, e.g., stationary point, travel, banding; distance travelled in case of travelling protocol); starting time of the sampling event; number of observers.

We used the eBird dataset released in December 2018^46^, focusing on records from 2005 to 2018, as data collected prior to 2005 were too scarce for analysis. We filtered this dataset to obtain high-quality checklists comparable in protocol and effort: we selected complete checklists only (i.e., in which observers explicitly declare having reported all bird species detected and identified); following either the ‘stationary points’ or the ‘travelling counts’ protocol; with durations of continuous observation of 0.5-10 hrs; with observers travelling distances during the checklist < 5 km; only from experienced observers (≥ 10 checklists; ≥ 30 species per checklist on average; ≥ 100 different species in total); and removing potentially duplicates (checklists with same day at same place).

After data filtering (more details in Supplementary Methods 1), we obtained the final dataset used in the analyses, consisting of 66,777 checklists, covering 5,467 species, from 6,838 observers, in eight hotspots (Extended Fig. 3–4; Extended Table 2).

### Site characteristics

Our analyses include two types of sites: checklist sites, corresponding to the coordinates of each eBird checklist analysed (used in analyses I and III); and background sites, corresponding to the centre points of a regular grid of 2×2 km covering evenly the whole area of each hotspot (used in analysis II). We characterised each site according to five characteristics – calculated in a 1-km radius buffer around its coordinates – two binary and three continuous: protected (if coordinates fall within a protected area^47^; Extended Fig. 5) versus non-protected; forest (if > 60% of the 1 km buffer around the point is forested^48^) versus non-forest (< 10% forested; sites with intermediate forest cover were removed from analyses); altitude^49^; agricultural suitability^50^; and remoteness^51^. In addition, we classified each forest site according to three continuous variables: canopy height^52^; forest contiguity (proportion of forest cover^48^, 0.6 to 1); and wilderness level (opposite of human footprint^53^).

Finally, checklist sites were also characterised according to four measures of local bird diversity: overall species richness (total number of species detected in the checklist); richness in forest-dependent species (high or medium dependency on forest habitats^31^); richness in endemic species (at least 90% of their global distribution within a hotspot^54^); and richness in species of concern (classified as Near Threatened or threatened, i.e., Vulnerable, Endangered, or Critically Endangered^31^; more details in Supplementary Methods 2).

### Index of observer expertise

Heterogeneity in observers’ birding skills increases data variability and potentially introduces biases to the analyses^55,56^. Heterogeneity is particularly high in citizen science datasets like eBird, where volunteers range from those only familiar with a few common local birds to experienced observers capable of detecting rare and cryptic species. As stated above, we only included checklists from relatively experienced observers. To account for the remaining variability in observer expertise, we calculated an observer expertise score (used as an explanatory variable in the statistical analyses), adapted from Kelling et al.^57^ and from Johnston et al.^56^, and calculated separately for each continent. It estimates the variation in the number of species that observers are predicted to detect in similar conditions. To do so, we first ran a mixed General Additive Model (function *gamm* from ‘mgcv’ R package^58^) modelling species richness of checklists against potential confounding variables that are expected to affect either the number of species detected (sampling protocol; n.observers number of observers; duration of sampling; time of the day) or the true species richness (lat latitude; lon longitude; and Julian day), adding observer as a random effect:

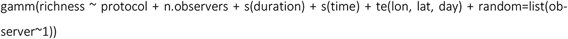

The notation s() indicates that the variable was used as a smoothed term; te() indicates that the variables have been used as interacting smooth terms, allowing here species richness to vary spatially during the year.

After fitting this model to each continental data subset, we used it to predict the logarithm of species richness that each observer would report for a fictive stationary point with all variables fixed to their median values. This resulted in an observer expertise score that we then assigned to all checklists; assigning the observer score of the observer with the highest expertise score in cases of multiple observers. This index ranged from 2.2 to 4.3 in Africa, from 2.3 to 4.4 in the Americas, and from 2.8 to 4.5 in Asia (more details in Supplementary Methods 3).

### Statistical analyses of protected area effectiveness

We investigated protected area effectiveness at retaining bird diversity through a set of three connected statistical analyses (Fig. 2), undertaken separately for each hotspot, using GAM models^58^. The first analysis (I) directly estimated the effects of protection on bird diversity while the two others (II and III) investigated the underlying mechanisms to explain the results of the first analysis.

Analysis I quantifies the effect of protected areas on bird diversity through models contrasting bird diversity of checklist sites between protected versus unprotected sites, while controlling for protected area location biases (and other potential confounding factors):

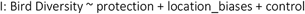

Analysis II quantifies the effectiveness of protected areas at mitigating forest loss and forest degradation, through models controlling for location biases and spatial autocorrelation. To measure the effects of protection on forest loss (IIa), we built logistic models contrasting protected versus unprotected background sites in their probability of being forested:

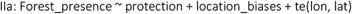

To measure the effects of protected areas on forest degradation (IIb), we built Gaussian models contrasting protected versus unprotected background forested sites in terms of forest quality (canopy height, forest contiguity, or wilderness):

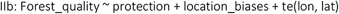

Analysis III quantifies the effects of forest presence (IIIa) or of forest quality (IIIb) on bird diversity, while controlling for potential confounding factors. In IIIa, we built models con-trasting bird diversity in forest versus non-forest checklist sites:

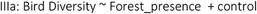

In IIIb, we modelled local bird diversity of forested sites against the three forest quality variables, as well as protected status in order to capture other aspects of forest quality that could be increased within protected areas (e.g. enforcement of hunting regulations; what we call protected area residuals):

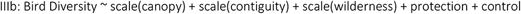

In analyses I and III, the response variable Bird Diversity is one of the four metrics of local bird diversity. We assumed Gaussian distribution for the overall richness, and a negative binomial distribution for the richness in forest-dependent species, endemic species and species of concern.

In analysis II, the response variable is either the binary Forest_presence (site forested or not) or each of three measures of Forest_quality (canopy height, forest contiguity, or wilderness).

The term location_bias in analyses I and II corresponds to s(altitude) + s(remoteness) + s(agricultural_suitability), supplemented by a control for spatial autocorrelation in analysis II with the term + te(lon, lat). It controls for potential biases in protected area location in relation to altitude, remoteness and agricultural suitability^16,17^ (Extended Fig. 6–8).

In analyses I and III, we controlled for other potential confounding factors that could affect the bird diversity reported in a checklist (Extended Fig. 9-16). In particular, we controlled for:heterogeneity in sampling effort (sampling duration; observer expertise; number of observers:n.observers); temporal effects (year to account for possible trends; day to account for season); spatial heterogeneity (lat latitude, lon longitude). The term control was thus:

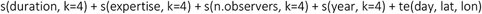

When the response variable was richness in forest-dependent species, in endemic species or in threatened and Near Threatened species, we also controlled for overall species richness, thus using as control term

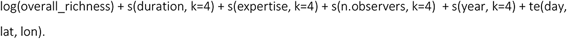

In analysis I, altitude is already controlled for under the location_bias term; in analysis III, the control term also includes a term controlling for it: s(altitude, k=6).

## Code availability

All R scripts will be deposited on an open repository after revision.

## Competing interests statement

The authors declare having no competing interests.

## References

1. UNEP-WCMC, IUCN and NGS. Protected Planet Report 2018. 70 (2018).

2. Watson, J. E. M., Dudley, N., Segan, D. B. & Hockings, M. The performance and potential of protected areas. Nature 515, 67–73 (2014).

3. Barlow, J. et al. The future of hyperdiverse tropical ecosystems. Nature 559, 517 (2018).

4. Lewis, S. L., Edwards, D. P. & Galbraith, D. Increasing human dominance of tropical forests. Science 349, 827–832 (2015).

5. Mittermeier, R. A. Hotspot revisited. (2004).

6. Cazalis, V., Belghali, S. & Rodrigues, A. S. L. Using a large-scale biodiversity monitoring dataset to test the effectiveness of protected areas at conserving North-American breeding birds. bioRxiv 433037, ver. 4peerreviewed and recommended by PCI Ecology (2019) doi:10.1101/433037.

7. Gray, C. L. et al. Local biodiversity is higher inside than outside terrestrial protected areas worldwide. Nature Communications 7, 12306 (2016).

8. Sullivan, B. L. et al. The eBird enterprise: An integrated approach to development and application of citizen science. Biological Conservation 169, 31–40 (2014).

9. Barnes, M. D., Glew, L., Wyborn, C. & Craigie, I. D. Prevent perverse outcomes from global protected area policy. Nature Ecology & Evolution 2, 759 (2018).

10. Visconti, P. et al. Protected area targets post-2020. Science eaav6886 (2019) doi:10.1126/science.aav6886.

11. Maxwell, S. L. et al. Area-Based Conservation in the 21st Century. Preprints 2020, 2020010104 (doi: 10.20944/preprints202001.0104.v1). (2020).

12. IPBES. Global assessment report on biodiversity and ecosystem services of the Intergovernmental SciencePolicy Platform on Biodiversity and Ecosystem Services. E. S. Brondizio, J. Settele, S. Díaz, and H. T. Ngo (editors). (2019).

13. SCBD. Aichi Biodiversity Targets. In: COP 10 Decision X/2: Strategic Plan for Biodiversity 2011–2020. (2010).

14. Pimm, S. L., Jenkins, C. N. & Li, B. V. How to protect half of Earth to ensure it protects sufficient biodiversity. Science Advances 4, eaat2616 (2018).

15. Pressey, R. L., Visconti, P. & Ferraro, P. J. Making parks make a difference: poor alignment of policy, planning and management with protected-area impact, and ways forward. Philosophical Transactions of the Royal Society B: Biological Sciences 370, 20140280 (2015).

16. Joppa, L. N. & Pfaff, A. High and far: biases in the location of protected areas. PLoS One 4, (2009).

17. Venter, O. et al. Bias in protected-area location and its effects on long-term aspirations of biodiversity conventions. Conservation Biology 32, 127–134 (2018).

18. Andam, K. S., Ferraro, P. J., Pfaff, A., Sanchez-Azofeifa, G. A. & Robalino, J. A. Measuring the effectiveness of protected area networks in reducing deforestation. Proceedings of the National Academy of Sciences 105, 16089–16094 (2008).

19. Barlow, J. et al. Anthropogenic disturbance in tropical forests can double biodiversity loss from deforestation. Nature 535, 144–147 (2016).

20. Newbold, T. et al. Ecological traits affect the response of tropical forest bird species to land-use intensity. Proceedings of the Royal Society of London B: Biological Sciences 280, 20122131 (2013).

21. Peres, C. A., Barlow, J. & Laurance, W. F. Detecting anthropogenic disturbance in tropical forests. Trends in Ecology & Evolution 21, 227–229 (2006).

22. Chandler, M. et al. Contribution of citizen science towards international biodiversity monitoring. Biological Conservation 213, 280–294 (2017).

23. Nelson, E. et al. Modeling multiple ecosystem services, biodiversity conservation, commodity production, and tradeoffs at landscape scales. Frontiers in Ecology and the Environment 7, 4–11 (2009).

24. Anderson, E. & Mammides, C. The role of protected areas in mitigating human impact in the world’s last wilderness areas. Ambio (2019) doi:10.1007/s13280-019-01213-x.

25. Geldmann, J., Manica, A., Burgess, N. D., Coad, L. & Balmford, A. A global-level assessment of the effectiveness of protected areas at resisting anthropogenic pressures. PNAS (2019) doi:10.1073/pnas.1908221116.

26. Coetzee, B. W. T., Gaston, K. J. & Chown, S. L. Local scale comparisons of biodiversity as a test for global protected area ecological performance: a meta-analysis. PLOS ONE 9, e105824 (2014).

27. Sullivan, B. L. et al. eBird: A citizen-based bird observation network in the biological sciences. Biological Conservation 142, 2282–2292 (2009).

28. Laurance, W. F. et al. Averting biodiversity collapse in tropical forest protected areas. Nature 489, 290–294 (2012).

29. Sorte, F. A. L. & Somveille, M. Survey completeness of a global citizen-science database of bird occurrence. Ecography 42, 1–10 (2019).

30. Olson, D. M. et al. Terrestrial Ecoregions of the World: A New Map of Life on EarthA new global map of terrestrial ecoregions provides an innovative tool for conserving biodiversity. BioScience 51, 933–938 (2001).

31. BirdLife International. IUCN Red List for birds. Version 2017.1. downloaded from <http://www.birdlife.org>. (2017).

32. Venter, O. et al. Sixteen years of change in the global terrestrial human footprint and implications for biodiversity conservation. Nature Communications 7, 12558 (2016).

33. Hillebrand, H. et al. Biodiversity change is uncoupled from species richness trends: Consequences for conservation and monitoring. Journal of Applied Ecology 55, 169–184 (2018).

34. Dornelas, M. et al. Assemblage time series reveal biodiversity change but not systematic loss. Science 344, 296–299 (2014).

35. Roxburgh, S. H., Shea, K. & Wilson, J. B. The intermediate disturbance hypothesis: patch dynamics and mechanisms of species coexistence. Ecology 85, 359–371 (2004).

36. Hiley, J. R., Bradbury, R. B. & Thomas, C. D. Impacts of habitat change and protected areas on alpha and beta diversity of Mexican birds. Diversity Distrib. 22, 1245–1254 (2016).

37. Eklund, J. et al. Contrasting spatial and temporal trends of protected area effectiveness in mitigating deforestation in Madagascar. Biological Conservation 203, 290–297 (2016).

38. Rutt, C. L., Jirinec, V., Cohn‐Haft, M., Laurance, W. F. & Stouffer, P. C. Avian ecological succession in the Amazon: A long-term case study following experimental deforestation. Ecology and Evolution 9, 13850–13861 (2019).

39. Bruner, A. G., Gullison, R. E., Rice, R. E. & Fonseca, G. A. B. Effectiveness of parks in protecting tropical biodiversity. Science 291, 125–128 (2001).

40. Giakoumi, S. & Pey, A. Assessing the Effects of Marine Protected Areas on Biological Invasions: A Global Review. Front. Mar. Sci. 4, (2017).

41. Hallmann, C. A. et al. More than 75 percent decline over 27 years in total flying insect biomass in protected areas. PLOS ONE 12, e0185809 (2017).

42. Craigie, I. D. et al. Large mammal population declines in Africa’s protected areas. Biological Conservation 143, 2221–2228 (2010).

43. Beaudrot, L. et al. Standardized Assessment of Biodiversity Trends in Tropical Forest Protected Areas: The End Is Not in Sight. PLOS Biology 14, e1002357 (2016).

44. Butchart, S. H. M. et al. Shortfalls and Solutions for Meeting National and Global Conservation Area Targets. Conservation Letters 8, 329–337 (2015).

45. Kroner, R. E. G. et al. The uncertain future of protected lands and waters. Science 364, 881–886 (2019).

## References

46. eBird. eBird: An online database of bird distribution and abundance [web application]. eBird, Cornell Lab of Ornithology, Ithaca, New York. Available: http://www.ebird.org. (Accessed: February 12, 2019, version Dec18). (2018).

47. UNEP-WCMC & IUCN. Protected Planet: [WDPA-shapefile-polygons; The World Database on Protected Areas (WDPA)/The Global Database on Protected Areas Management Effectiveness (GD-PAME)] [On-line, downloaded 02/10/2018], Cambridge, UK. <www.protectedplanet.net>. (2018).

48. ESA. Climate Change Initiative - Land cover project map v2.0.7. Data from year 2015. <http://maps.elie.ucl.ac.be/CCI/viewer/index.php>. (2015).

49. National Geophysical Data Center. Global Land One-kilometer Base Elevation (GLOBE), version 1. <https://www.ngdc.noaa.gov/mgg/topo/gltiles.html>. (1999) doi:10.7289/V52R3PMS.

50. Zabel, F., Putzenlechner, B. & Mauser, W. Global Agricultural Land Resources – A High Resolution Suitability Evaluation and Its Perspectives until 2100 under Climate Change Conditions. PLoS One 9, (2014).

51. Weiss, D. J. et al. A global map of travel time to cities to assess inequalities in accessibility in 2015. Nature 553, 333–336 (2018).

52. Simard, M., Pinto, N., Fisher, J. B. & Baccini, A. Mapping forest canopy height globally with spaceborne lidar. Journal of Geophysical Research: Biogeosciences 116, (2011).

53. Venter, O. et al. Global terrestrial Human Footprint maps for 1993 and 2009. Scientific Data 3, 160067 (2016).

54. BirdLife International and HBW. Bird species distribution maps of the world. Version 7.0. Available at <http://datazone.birdlife.org/species/requestdis>. (2017).

55. Dickinson, J. L., Zuckerberg, B. & Bonter, D. N. Citizen Science as an Ecological Research Tool: Challenges and Benefits. Annual Review of Ecology, Evolution, and Systematics 41, 149–172 (2010).

56. Johnston, A., Fink, D., Hochachka, W. M. & Kelling, S. Estimates of observer expertise improve species distributions from citizen science data. Methods in Ecology and Evolution 9, 88–97 (2018).

57. Kelling, S. et al. Can Observation Skills of Citizen Scientists Be Estimated Using Species Accumulation Curves? PLOS ONE 10, e0139600 (2015).

58. Wood, S. N. Fast stable restricted maximum likelihood and marginal likelihood estimation of semiparametric generalized linear models: Estimation of Semiparametric Generalized Linear Models. Journal of the Royal Statistical Society: Series B (Statistical Methodology) 73, 3–36 (2011).

