## Supplementary methods for "Effectiveness of protected areas in conserving tropical forest birds"

### 1. Data selection: eBird checklists

#### 1A. Spatial filtering

We focused on global biodiversity hotspots (Mittermeier, 2004) which overlapped by more than 25% of their extent the “tropical and subtropical moist broadleaf forests” biome (boundaries from Olson et al. (2001)). We obtained 16 hotspots: Atlantic Forest, Tropical Andes, Tumbes-Chocó-Magdalena, Caribbean Islands, Mesoamerica, Guinean Forests of West Africa, Eastern Afromontane, Coastal Forests of Eastern Africa, Madagascar, Western Ghats and Sri Lanka, Indo-Burma, Sundaland, Philippines, East Melanesian Islands, and New Caledonia.

Of these, we ultimately analysed only eight hotspots for which we obtained more than 1,000 eBird checklists (after following the data filtering procedure detailed below). We first downloaded the eBird checklists for the respective countries per hotspot (eBird codes between brackets):

- **Atlantic Forest**: Argentina [AR], Brazil [BR], Paraguay [PY].
- **Tropical Andes**: Argentina [AR], Bolivia [BO], Colombia [CO], Ecuador [EC], Peru [PE], Venezuela [VE].
- **Tumbes-Choco-Magdalena**: Colombia [CO], Ecuador [EC], Panama [PA], Peru [PE].
- **Mesoamerica**: Belize [BZ], Costa Rica [CR], Guatemala [GT], Honduras [HN], Mexico [MX], Nicaragua [NI], Panama [PA], El Salvador [SV].
- **Eastern Afromontane**: Burundi [BI], Democratic Republic of Congo [CD], Eritrea [ER], Ethiopia [ET], Kenya [KE], Mozambique [MZ], Rwanda [RW], Sudan [SD], South Sudan [SS], Tanzania [TZ], Uganda [UG].
- **Western Ghats and Sri Lanka**: India [IN], Sri Lanka [LK].
- **Indo-Burma**: Bangladesh [BD], Hong Kong [HK], India [IN], Cambodia [KH], Laos [LA], Myanmar [MM], Malaysia [MY], Thailand [TH], Vietnam [VN], China [CN].
- **Sundaland**: Brunei [BN], Indonesia [ID], Malaysia [MY], Thailand [TW].

We then filtered the checklists to include only those overlapping both the hotspot and the “tropical and subtropical moist broadleaf forests” biome.

In the Indo-Burma hotspot, we excluded records from China [CN] (19% of the hotspot, 2% of checklists) because no protected area data were available for this country in the publicly available version of the World Database on Protected Areas (UNEP-WCMC and IUCN, 2018).

#### 1B. Filtering by sampling protocol

In order to be able to treat this dataset as presence/absence, we focused on checklists for which observers stated that they reported every species detected (Sullivan et al., 2009). Accordingly, we also removed checklists unlikely to capture all species because using particular protocols (e.g. banding) or targeting specific groups (e.g. waders, nocturnal). We therefore used only checklists for which the protocol reported was either ‘stationary points’ or ‘travelling counts’. In stationary points, observers remain at the checklist location, and report both the starting time and the duration of the sampling. In travelling counts, moving observers report the checklist location (usually the mid-point of their itinerary), starting time, duration of the sampling and distance travelled. We excluded travelling counts with a travel distance >5 km, as they may not represent the local bird composition around the reported GPS location. To further increase comparability, we excluded sampling events that were very short (<30 minutes) or very long (>10 hours).

We also included some data classified in the eBird dataset under the protocol category ‘historical counts’, which consist of sampling events for which birding was the primary focus but for which the observer was not able to fill all fields required for reporting stationary points or travel counts (e.g. starting time, duration, distance). We used historical counts if duration was known and ranged from 30 minutes to 10 hours, and if distance was known and shorter than 5 km.

#### 1C. Observation filtering

We excluded observations that were disapproved by the eBird review process, corresponding to exotic, feral or escaped individuals. Established introduced species were kept in the dataset.

Using the *auk_rollup* function from the above-mentioned R package ‘auk’ (Strimas-Mackey et al., 2017), we brought all observations of subspecies to the species level.

#### 1D. Filtering checklists based on observer experience

To aim for complete checklists, we filtered observations to retain only those submitted by relatively experienced observers. In order to identify these, we analysed checklists per observer across each of three continents, defined according to the following list of countries (eBird codes between brackets):

- **Americas**: Antigua and Barbuda [AG], Anguilla [AI], Argentina [AR], Bolivia [BO], Brazil [BR], Bahamas [BS], Belize [BZ], Chile [CL], Colombia [CO], Costa Rica [CR], Cuba [CU], Dominica [DM], Dominican Republic [DO], Ecuador [EC] Falkland Islands [FK], Grenada [GD], French Guyana [GF], Guadeloupe [GP], Guatemala [GT], Guyana [GY], Honduras [HN], Haiti [HT], Jamaica [JM], Saint Knitts and Nevis [KN], Cayman Islands [KY], Saint Lucia [LC], Martinique [MQ], Montserrat [MS], Mexico [MX], Nicaragua [NI], Panama [PA], Peru [PE], Puerto Rico [PR], Paraguay [PY], Suriname [SR], El Savador [SV], Turks and Caicos Islands [TC], Uruguay [UY], Saint Vincent and the Grenadines [VC], Venezuela [VE], British Virgin Islands [VG], Virgin Islands [VI].
- **Asia**: Bangladesh [BD], Brunei [BN], Bhutan [BT], China [CN], Hong Kong [HK], Indonesia [ID], India [IN], Cambodia [KH], Laos [LA], Sri Lanka [LK], Myanmar [MM], Malaysia [MY], Nepal [NP], Papua New Guinea [PG], Philippines [PH], Pakistan [PK], Thailand [TH], Taiwan [TW], Vietnam [VN].
- **Africa**: Angola [AO], Burkina Faso [BF], Burundi [BI], Benin [BJ], Botswana [BW], Democratic Republic of Congo [CD], Central African Republic [CF], Congo [CG], Cote d’Ivoire [CI], Cameroon [CM], Djibouti [DJ], Algeria [DZ], Egypt [EG], Western Sahara [EH], Eritrea [ER], Ethiopia [ET], Gabon [GA], Ghana [GH], Gambia [GM], Guinea [GN], Equatorial Guinea [GQ], Guinea-Bissau [GW], Kenya [KE], Liberia [LR], Lesotho [LS], Libya [LY], Morocco [MA], Madagascar [MG], Mali [ML], Mauritania [MR], Malawi [MW], Mozambique [MZ], Namibia [NA], Niger [NE], Nigeria [NG], Rwanda [RW], Sudan [SD], Sierra Leone [SL], Senegal [SN], Somalia [SO], South Sudan [SS], Sao Tomé and Principe [ST], Swaziland [SZ], Chad [TD], Togo [TG], Tunisia [TN], Tanzania [TZ], Uganda [UG], South Africa [ZA], Zambia [ZM], Zimbabwe [ZW].

Within each given continent, we defined as ‘experienced observers’ those who had submitted ≥ 10 checklists to eBird, with ≥ 30 species per checklist on average, and covering ≥ 100 different species in total. We only retained the checklists by the observers who were classified as ‘experienced’ in the corresponding continent.

#### 1E. Removing duplicates

Multiple observations of the same birds can happen either because several observers travelled together or because they came independently to the same site on the same day, both situations creating pseudo-replication.

When submitting checklists, observers can specify if they were observing with others. For checklists in this situation, we removed duplication by keeping only one checklist (the one with the lowest checklist id code), using the *auk_unique* function implemented in the R package ‘auk’ (a package specifically created to process eBird data; (Strimas-Mackey et al., 2017)).

In addition, we filtered for other possible duplicates by independent travellers: whenever two checklists with equal dates were reported with less than 2 km between them, we randomly selected one of them.

#### 1F. Taxonomic standardisation

The taxonomic classification used in eBird follows the Clements taxonomy (Clements et al., 2018). In order to be able to cross the bird observation dataset with the species’ trait data (section 2 below) we have converted it to the taxonomy used by BirdLife International and HBW (2017). For this, we used an unpublished table kindly provided by the Cornell Lab of Ornithology, which summarises relationships between the two taxonomies, by applying the following rules:

- In the case of a simple difference in name, we applied the Birdlife name to the eBird records (295 species).
- Whenever a single species in the BirdLife list was treated as multiple species in eBird, we lumped the eBird records (92 species).
- Whenever multiple species in the BirdLife list were treated as a single species in eBird, we split the eBird records based on the BirdLife distribution maps for the corresponding species (358 species). Any records outside the BirdLife distribution maps were assigned to the species whose distribution was the closest. In the extreme rare case of overlap between distributions of these species (12 over the 5,467 species for a total of ~1,500 observations), observations falling within the distribution overlap were all assigned to a single of the two species (selected randomly between both).

The lists of species in each case are detailed in the supplementary information spreadsheet “Species taxonomy changes supplementary table”.

#### 1G. Final dataset analysed

After the above steps, plus the removal of sites of intermediate forest cover (see section 2A below), we obtained a total of 66,777 checklists, covering 5,467 species, from 6,838 observers, in eight hotspots. This was the final dataset used in the analyses. For further details, and a breakdown per hotspot, see Extended Table 2 and Extended Fig. 3-4.

### 2. Site characteristics

Our analyses include two types of sites: checklist sites, corresponding to the coordinates of each eBird checklist analysed (used in analyses I and III, see below); and background sites, corresponding to the centre points of a regular grid of 2×2 km covering evenly the whole area of each hotspot (used in analysis II).

We characterised each site according to five variables: two binary (protected vs. non-protected; forest vs. non-forest) and three continuous (altitude; agricultural suitability; remoteness). For forest sites, we characterised them according to three additional continuous variables (canopy height; forest contiguity; and wilderness level).

Checklist sites were also characterised according to four measures of local bird diversity (richness of all species, of forest-dependent species, of endemic species, and of threatened and Near Threatened species).

#### 2A. Protection (binary)

A site was considered ‘protected’ if its coordinates overlapped a protected area, as mapped in the World Database on Protected Areas (UNEP-WCMC and IUCN, 2018). As is the standard protocol in global analysis of protected area coverage (UNEP-WCMC and IUCN, 2019), we excluded: “Man and Biosphere” reserves; protected areas without associated polygons; protected areas that did not have as status “designated”, “inscribed” or “established”.

We would have obtained similar results if we had instead derived the protection status from the proportion of area under protection within a 1-km buffer around the site, as the vast majority of buffers are protected by either 0 or 100 % (Extended Fig. 5).

#### 2A. Forest habitat (binary)

To derive whether a site was forested or not, we used the 2015 version of Climate Change Initiative Land Cover layer, with a resolution of 300 m (ESA, 2015). We considered as forests all categories described as strict Tree Cover (i.e. codes 50-90, 160 and 170), and as non-forest all others (but excluding from the analyses water bodies, code 210).

For each site, we first calculated the percentage of pixels overlapping the 1-km buffer that were forest. We then classified as ‘forest’ the sites with >60% forest, and as ‘non-forest’ those with <10% forest. We thus obtained two types of localities very contrasting in their forest cover, removing from the analyses all sites with intermediate (10 to 60%) cover.

#### 2C. Altitude (continuous)

Altitude data were obtained from the GLOBE Digital Elevation Model (National Geophysical Data Center, 1999), which has a 0.008 degree resolution (~930 m at latitude 0). We calculated the altitude per site as the median of the values intersecting a 1-km buffer around the site.

#### 2D. Agricultural suitability (continuous)

We used a global raster of resolution ~1-km mapping a value of agricultural suitability (Zabel et al., 2014). Their model estimates for each cell the suitability of each of the 16 most important food and energy crops in the world – based on climatic conditions, soil and topography – and assigns to the cell the value of the crop with the highest suitability. It has no unit and is included in a 0 – 100 interval. We obtained a value of agricultural suitability per site as the median of the values intersecting a 1-km buffer around the site.

#### 2E. Remoteness (continuous)

Remoteness was derived from the global accessibility map of resolution ~1-km, which estimates the travel time needed for a human to reach the nearest city with ≥ 1,500 inhabitants (Weiss et al., 2018). We obtained a remoteness value per site as the median of the values intersecting a 1-km buffer around the site.

#### 2F. Canopy height (continuous; forest sites only)

We used a global raster of canopy height at resolution ~1-km derived from spaceborne light detection and ranging (lidar) data (Simard et al., 2011). We calculated the canopy height value for all forest sites as the median of the values intersecting a 1-km buffer around the site.

#### 2G. Forest contiguity (continuous; forest sites only)

Using the above-mentioned forest layer (used to classify sites as forest or not), we assigned to each forest site the proportion of forest cover (0.6 to 1) as an index of forest contiguity.

#### 2H. Wilderness level (continuous; forest sites only)

We used the 2009 global terrestrial human footprint map (Venter et al., 2016), with a resolution ~1-km, obtained by combining spatial information on human pressures including human infrastructures, agricultural land use and population density. For each forest site, we obtained a wilderness value as the opposite of the median human footprint (-1 * human footprint) across pixels intersecting the 1-km buffer around the site.

#### 2I. Overall species richness

For each checklist site, we calculated the total number of species detected in the checklist.

#### 2H. Richness in forest-dependent species

For each checklist site, we calculated the total number of species classified as ‘forest-dependent’. These are species classified as either highly- or medium-dependent in a pre-existing classification by BirdLife International (2017) that includes five categories:

- Highly-dependent**:** Forest specialists; characteristic of the interior of undisturbed forest; may persist in secondary forest and forest patches if their particular ecological requirements are met, but where they do occur away from the interior, they are usually less common; rarely seen in non-forest habitats; breeding is almost invariably within forest.
- Medium-dependent**:** Forest generalists; may occur in undisturbed forest but also regularly found in forest strips, edges and gaps; likely to be commoner in such situations and in secondary forest than in the interior of intact forest; breeding is typically within forest.
- Low-dependent**:** Often recorded in forest, but not dependent on it; almost always more common in non-forest habitats, where most likely to breed.
- Non-forest species**:** Does not normally occur in forest.
- Unknown (none in the dataset): Occurs or probably occurs in forest, but dependency on it is unknown, but could be high.

#### 2J. Richness in endemic species

For each checklist site, we calculated the total number of species that are classified as ‘endemic’ to the corresponding hotspot. This includes all species with at least 90% of its global distribution (BirdLife International and HBW, 2017) contained within the boundary of the hotspot (considering the whole area of the hotspot, not only the part included in the “tropical and subtropical moist broadleaf forests” biome).

#### 2K. Richness in threatened and Near Threatened species

For each checklist site, we calculated the total number of species classified as being either threatened (Vulnerable, Endangered, or Critically Endangered) or Near Threatened in the International Union for Conservation of Nature (IUCN) Red List of Threatened Species (BirdLife International, 2017).

### 3. Index of observer expertise

For each observer in our analysis, we derived an expertise score using an index adapted from Kelling et al. (2015) and from Johnston et al. (2018). Calculated separately for each continent, the index estimates the variation in the number of species that observers are predicted to detect in similar conditions.

We first ran a mixed General Additive Model (function *gamm* from ‘mgcv’ R package (Wood, 2011)) modelling the species richness of checklists against several sampling variables that are expected to affect species richness, adding observer (i.e., observer individual identifying number) as a random effect:

gamm(richness ~ protocol + n.observers + s(duration) + s(time) + te(lon, lat, day) + random=list(observer~1))

To control for differences in checklist sampling effort, we included both duration (of sampling, in minutes) and time (starting hour of sampling) as smooth terms, allowing non-linear correlations. We included lon (longitude, in decimal degrees), lat (latitude, in decimal degrees) and day (Julian date, from 1 to 365/366) as a smoothed three-way interaction, thus allowing richness to vary across space and season of the year. We opted for a GAM (following Kelling et al. (2015)) rather than a Generalised Linear Model (as Johnston et al. (2018)) because the former allows for non-linear effects. For fitting issues, we only included a random effect on intercept, rather than also on the slope between duration and richness, as in Kelling et al. (2015).

Following Johnston et al. (2018), we fitted this model to a nearly complete dataset per continent (defined as in section 1D), i.e., before the filtering steps detailed in sections 1 and 2A. The only filtering rules applied were to exclude: observations prior to 2005; disapproved observations; checklists that did not report all species observed (because the model is based on richness). We also removed checklists that did not report one of the covariates (about 15% of observations), given that the GAMM cannot accommodate empty records.

We assumed that species richness followed a Poisson distribution, as in Kelling et al. (2015) and Johnston et al. (2018), because the dataset includes many checklists with low richness, making the distribution closer to a Poisson than a Gaussian distribution.

Having fitted the model to the data, we then used it to predict the species richness that each observer would report for a fictive stationary point with all variables fixed to their median values. The observer expertise score, measured as the logarithm of this predicted species richness, ranged from 2.2 to 4.3 in Africa, from 2.3 to 4.4 in the Americas, and from 2.8 to 4.5 in Asia. We then assigned to each checklist used in our analysis the expertise score of the observer. In case of multiple observers, we assigned the score of the observer with the highest expertise score. This score was then used as an explanatory variable in the statistical analyses below.

### 2. Statistical analyses of protected area effectiveness

We investigated protected area effectiveness at retaining bird diversity through a set of three connected statistical analyses, undertaken separately for each hotspot (Fig. 2). We used General Additive Models (GAMs) for all analyses, which are similar to Generalised Linear Models, but accommodate nonlinear relationships between response and explanatory variables (Wood, 2011; Zuur et al., 2009).

We implemented these models using the ‘mgcv’ R package (Wood, 2011), running an independent model for each hotspot and for each response variable.

In analyses I and III, we assumed a negative binomial distribution for all models, except for those where overall species richness was the response variable. For the later, the distribution obtained after the filtering of checklists was closer to a Gaussian distribution, so we assumed that instead. In analysis II, we assumed a Binomial distribution for forest presence and Gaussian distributions for the three variables of forest quality.

#### 4A. Analysis I: effect of protected areas on bird diversity

Analysis I quantifies the effect of protected areas on the bird diversity reported in checklists, controlling for site location biases and other potential confounding factors. The model has the following structure:

Bird Diversity ~ protection + location_biases + control

where:

- Bird Diversity corresponds to one of the four bird diversity indices: overall richness; richness in forest-dependent species; richness in endemics; richness in threatened and Near Threatened species;
- Protection corresponds to the binary variable indicating whether the site is protected (1) or not (0);
- location_biases corresponds to a term controlling for eventual spatial biases on the location of protected areas, formalised as: s(altitude) + s(remoteness) + s(agricultural_suitability), corresponding to the use of these three variables in independent smoothed terms (allowing non-linear relationships) without limiting the curves complexity (Extended Fig. 7);
- control corresponds to a term accounting for heterogeneity in sampling effort and potential spatiotemporal variation in bird diversity metrics. In models using the overall species richness as response variable, this was formalised as: s(duration, k=4) + s(expertise, k=4) + s(year, k=4) + te(day, lat, lon). For all other models, we also controlled for overall species richness, so it became: log(overall_richness) + s(duration, k=4) + s(expertise, k=4) + s(n.observers, k=4) + s(year, k=4) + te(lat, lon, day). Where:
  - s(duration, k=4) is the sampling duration in minutes, used here as an independent smoothed term with the degree of the smoothing function fixed to 4, in order to limit the curve complexity (Extended Fig. 9-16);
  - s(expertise, k=4) is the observer expertise score for the checklist, used here as an independent smoothed term with the degree of the smoothing function fixed to 4, in order to limit the curve complexity (Extended Fig. 9-16);
  - s(n.observers, k=4) is the number of observers present during the sampling
  - s(year, k=4) is the year of the observation, included to account for potential temporal trends in the region, used here as an independent smoothed term with the degree of the smoothing function fixed to 4, in order to limit the curve complexity (Extended Fig. 9-16);
  - te(lat, lon, day) are the site’s decimal coordinates and the Julian date of the observation (from 1 to 365/366), used as a three-way interaction smoothed-term, allowing bird diversity indices to vary spatially during the year (e.g. a species can occur in a region more often during the winter while occurring more often in another region during the summer), thus enabling to account for migration patterns (Extended Fig. 9-16);
  - log(overall_richness) is the logarithm of the overall species richness. It was used in all models with richness in forest-dependent species, endemic species, and threatened and Near Threatened species as response variable (used in log because we assumed Negative Binomial distributions for these three variables). Therefore, these models test the effect of protection or habitat on the richness in forest-dependent species, endemic species or threatened and Near Threatened, for a given overall species richness (Extended Fig. 9-16).

#### 4B. Analysis II: effect of protected areas on forest quantity and quality

Analysis II quantifies the effect of protected areas at mitigating forest loss (analysis IIa) or forest degradation (analysis IIb), controlling for site location biases. Model were built from all background sites within each hotspot.

The model structure for analysis IIa is:

Forest_presence ~ protection + location_biases + te(lon, lat)

where:

- Forest_presence corresponds to the binary variable indicating whether the site is forested (1) or not (0);
- Protection corresponds to the binary variable indicating whether the site is protected (1) or not (0);
- location_biases corresponds to a term controlling for the possibility of location biases in protected areas (as above) (Extended Fig. 8);
- te(lon, lat) corresponds to a two-way interaction smoothed term, used here to control for spatial autocorrelation in habitat variables.

The models for analysis IIb were restricted to forest sites and have the following structure:

Forest_quality ~ protection + location_biases + te(lon, lat)

where:

- Forest_quality corresponds to one of the three continuous variable used for forest quality: canopy height, forest contiguity, and wilderness;
- Protection corresponds to the binary variable indicating whether the site is protected (1) or not (0);
- location_biases corresponds to a term controlling for the possibility of location biases in protected areas (as above) (Extended Fig. 8);
- te(lon, lat) corresponds to a two-way interaction smoothed term, used here to control for spatial autocorrelation in habitat variables.

#### 4C. Analysis III: effect of forest presence and quality on bird diversity

Analysis III quantifies the effects of forest presence (IIIa) and forest quality (IIIb) on the bird diversity reported in checklists.

The model structure for analysis IIIa is:

Bird Diversity ~ Forest_presence + control

where:

- Bird Diversity corresponds to one of the four bird diversity indices: overall richness; richness in forest-dependent species; richness in endemics; richness in threatened and Near Threatened species;
- control corresponds to a term accounting for heterogeneity in sampling effort and potential spatiotemporal variation in bird diversity metrics. It corresponds to the term used in analysis I, but supplemented with s(altitude, k=6), the altitude in meters, used here as an independent smoothed term with the degree of the smoothing function fixed to 4, in order to limit the curve complexity;

The models for analysis IIIb were restricted to forest sites and have the following structure:

Bird Diversity ~ scale(canopy) + scale(contiguity) + scale(wilderness) + protection + control

where:

- Bird Diversity corresponds to one of the four bird diversity indices: overall richness; richness in forest-dependent species; richness in endemics; richness in threatened and Near Threatened species;
- canopy, contiguity, and wilderness respectively the canopy height, forest contiguity, and wilderness of checklist sites;
- scale() indicates that the variable has been scaled (by subtracting the mean and dividing by the standard deviation), so that their effect size are comparable;
- Protection corresponds to the binary variable indicating whether the site is protected (1) or not (0);
- control corresponds to a term accounting for heterogeneity in sampling effort and potential spatiotemporal variation in bird diversity metrics. It corresponds to the term used in analysis I, but supplemented with s(altitude, k=6), the altitude in meters, used here as an independent smoothed term with the degree of the smoothing function fixed to 4, in order to limit the curve complexity;

#### 4D. Potential effects of differences in habitat between protected and non-protected sites

In analyses I and IIa, we contrasted protected versus unprotected sites in order to investigate the effects of protection on either bird diversity or on the presence of forest. For this to be a perfect counterfactual analysis, the contrasts ought to have controlled for any confounding effects that make the sites distinct in other ways besides protection.

One possible confounding effect is that protected versus non-protected sites may not have had the same initial habitat, here assumed to be forest. If so, and if protection is then biased towards a particular habitat (e.g., forest) then the effects observed in analysis I and IIa (Figs. 3, 4) may also reflect either intrinsic or pre-existing differences in habitat rather than the effect of protection itself. Intrinsic differences refer to different natural vegetation (e.g. *páramo* vegetation in the high Andes), in which case the observed contrast between protected versus unprotected sites is affected by natural differences in community composition. Pre-existing differences correspond to habitat differences in place at the time of protection (e.g. if protected areas were towards remaining vegetation patches), in which case the contrasts observed may reflect the history of habitat change (e.g. deforestation) prior to protection rather than the effects of protection alone.

We aimed to reduce intrinsic habitat differences by focusing analysis within the “tropical and subtropical moist broadleaf forests” biome. According to Nelson and Chomitz (2009), this biome “contains the maximum extent of the world’s tropical and subtropical moist broadleaf forests” but it is unlikely that it would have been 100% forested across all of our study area. A global map of potential vegetation (Ramankutty and Foley (1999); raster at a resolution of 5’’, ~ 9 km at the Equator), representing the “vegetation that would most likely exist now in the absence of human activities”, predicts that for five out of eight hotspots the area we considered in these analyses would have been originally covered by forests by >80%: Mesoamerica 87%; Tumbes-Chocó-Magdalena 92%; Western Ghats and Sri Lanka 83%; Indo-Burma 90%; Sundaland 96% (considering as forest in the Ramankutty and Foley map: “Tropical Evergreen Woodland”, “Tropical Deciduous Woodland”, “Temperate Evergreen Woodland”, “Temperate Deciduous Woodland”, “Mixed Woodland”). For three other hotspots, it estimates lower proportions of original forest: Atlantic Forest 69%, Tropical Andes 67%, Eastern Afromontane 30%. Ramankutty and Foley (2009)’s method is however likely to underestimate original forest cover by discounting old forest conversion. For the Atlantic Forest, we also analysed a different map of predicted original vegetation produced by the Brazil Institute of Geography and Statistics, according to which our study area was 90% originally covered forest (IBGE - EMBRAPA, 2001). We were unable to find another reconstruction of original habitat for the whole of the Eastern Afromontane region, but a study focusing on the Eastern Arc Mountain (south-east part of the hotspot) mapped the original forest using paleoecological data and found that it was mostly covered by forest (Hall et al., 2009). In the Tropical Andes, much of the disagreement with Ramankutty and Foley (2009) corresponds to high altitude grasslands (*páramo*) that were indeed probably not forested. In summary, then, the study areas in this study were likely dominated by forest, but not uniformly.

We were unable to account directly for pre-existing habitat differences, given that many protected areas were designated prior to the availability of forest cover maps.

Nevertheless, the control covariates used in analyses I and IIa to account for location biases are likely to control to some extent for intrinsic as well as pre-existing habitat differences. Altitude affects both of these, as it affects both natural habitat (e.g. the transition from cloud forest to *páramo* in the Tropical Andes) and the likelihood, and thus history, of habitat conversion. Agricultural suitability is a factor of habitat conversion (Joppa and Pfaff, 2009), but models of agricultural suitability are generated from variables such as climate (rain, temperature, radiation), soil, and terrain (slope, exposure) that also affect the natural habitat of each site.

In summary then, we expect intrinsic habitat differences between protected and non-protected sites to have been relatively minor in this study, and that both these and pre-existing differences have been adequately controlled for in the statistical models.

### Supplementary references

BirdLife International (2017). IUCN Red List for birds. Version 2017.1. downloaded from <http://www.birdlife.org>.

BirdLife International and HBW (2017). Bird species distribution maps of the world. Version 7.0. Available at <http://datazone.birdlife.org/species/requestdis>.

Clements, J.F., Schulenberg, T.S., Iliff, M.J., Roberson, D., Fredericks, T.A., Sullivan, B.L., and Wood, C.L. (2018). The eBird/Clements checklist of birds of the world: v2018 [Downloaded from <http://www.birds.cornell.edu/clementschecklist/download/>].

ESA (2015). Climate Change Initiative - Land cover project map v2.0.7. Data from year 2015. <http://maps.elie.ucl.ac.be/CCI/viewer/index.php>.

Hall, J., Burgess, N.D., Lovett, J., Mbilinyi, B., and Gereau, R.E. (2009). Conservation implications of deforestation across an elevational gradient in the Eastern Arc Mountains, Tanzania. Biol. Conserv. *142*, 2510–2521.

IBGE - EMBRAPA (2001). Mapa de Solos do Brasil. Rio de Janeiro - Escala 1:5.000.000. The original vegetation map (shapefile format) can be downloaded for Brazil or Brazilian Legal Amazon limits.

Johnston, A., Fink, D., Hochachka, W.M., and Kelling, S. (2018). Estimates of observer expertise improve species distributions from citizen science data. Methods Ecol. Evol. *9*, 88–97.

Joppa, L.N., and Pfaff, A. (2009). High and far: biases in the location of protected areas. PLOS ONE *4*.

Kelling, S., Johnston, A., Hochachka, W.M., Iliff, M., Fink, D., Gerbracht, J., Lagoze, C., La Sorte, F.A., Moore, T., Wiggins, A., et al. (2015). Can Observation Skills of Citizen Scientists Be Estimated Using Species Accumulation Curves? PLOS ONE *10*, e0139600.

Mittermeier, R.A. (2004). Hotspot revisited (Cemex).

National Geophysical Data Center (1999). Global Land One-kilometer Base Elevation (GLOBE), version 1. <https://www.ngdc.noaa.gov/mgg/topo/gltiles.html>.

Nelson, A., and Chomitz, K.M. (2009). Protected area effectiveness in reducing tropical deforestation (Washington DC, USA: Independent Evaluation Group, The World Bank).

Olson, D.M., Dinerstein, E., Wikramanayake, E.D., Burgess, N.D., Powell, G.V.N., Underwood, E.C., D’amico, J.A., Itoua, I., Strand, H.E., Morrison, J.C., et al. (2001). Terrestrial Ecoregions of the World: A New Map of Life on EarthA new global map of terrestrial ecoregions provides an innovative tool for conserving biodiversity. BioScience *51*, 933–938.

Ramankutty, N., and Foley, J.A. (1999). Estimating historical changes in global land cover: Croplands from 1700 to 1992. Glob. Biogeochem. Cycles *13*, 997–1027.

Simard, M., Pinto, N., Fisher, J.B., and Baccini, A. (2011). Mapping forest canopy height globally with spaceborne lidar. J. Geophys. Res. Biogeosciences *116*.

Strimas-Mackey, M., Miller, E., and Hochachka, W. (2017). auk: eBird Data Extraction and Processing with AWK.

Sullivan, B.L., Wood, C.L., Iliff, M.J., Bonney, R.E., Fink, D., and Kelling, S. (2009). eBird: A citizen-based bird observation network in the biological sciences. Biol. Conserv. *142*, 2282–2292.

UNEP-WCMC, and IUCN (2018). Protected Planet: [WDPA-shapefile-polygons; The World Database on Protected Areas (WDPA)/The Global Database on Protected Areas Management Effectiveness (GD-PAME)] [On-line, downloaded 02/10/2018], Cambridge, UK. <www.protectedplanet.net>.

UNEP-WCMC, and IUCN (2019). Calculating protected area coverage. [On-line, consulted 06/02/2019]. <www.protectedplanet.net/c/calculating-protected-area-coverage>.

Venter, O., Sanderson, E.W., Magrach, A., Allan, J.R., Beher, J., Jones, K.R., Possingham, H.P., Laurance, W.F., Wood, P., Fekete, B.M., et al. (2016). Global terrestrial Human Footprint maps for 1993 and 2009. Sci. Data *3*, 160067.

Weiss, D.J., Nelson, A., Gibson, H.S., Temperley, W., Peedell, S., Lieber, A., Hancher, M., Poyart, E., Belchior, S., Fullman, N., et al. (2018). A global map of travel time to cities to assess inequalities in accessibility in 2015. Nature *553*, 333–336.

Wood, S.N. (2011). Fast stable restricted maximum likelihood and marginal likelihood estimation of semiparametric generalized linear models: Estimation of Semiparametric Generalized Linear Models. J. R. Stat. Soc. Ser. B Stat. Methodol. *73*, 3–36.

Zabel, F., Putzenlechner, B., and Mauser, W. (2014). Global Agricultural Land Resources – A High Resolution Suitability Evaluation and Its Perspectives until 2100 under Climate Change Conditions. PLoS ONE *9*.

Zuur, A.F., Ieno, E.N., Walker, N., Saveliev, A.A., and Smith, G.M. (2009). Mixed effects models and extensions in ecology with R (New York, NY: Springer).
