## Extended figures for "Effectiveness of protected areas in conserving tropical forest birds"

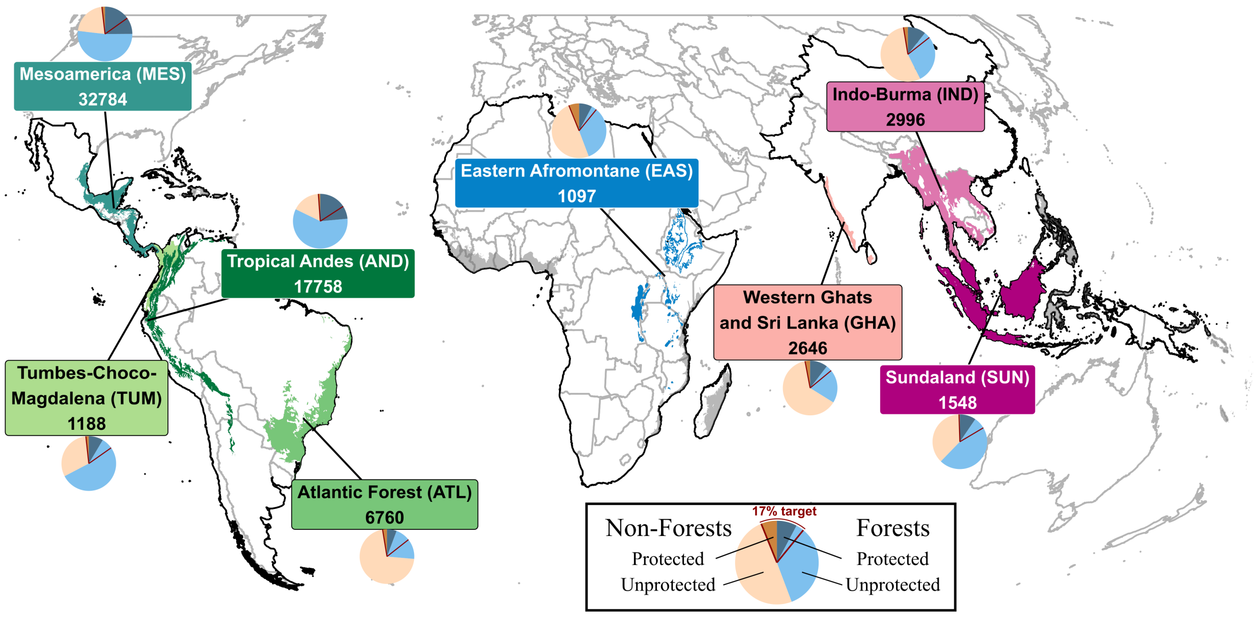


Extended Figure 1: Regions covered by the present study, with number of checklists and levels of protection for each hotspot. Coloured regions correspond to the area analysed within each of the eight hotspots, i.e. the intersection between the hotspot boundary and the "tropical and subtropical moist broadleaf forests" biome. For these, boxes indicate the hotspot name, acronym, and number of checklists analysed. Pie plots represent the proportion of forest/non-forest and protected/non-protected background sites in each hotspot. Red lines in pie plots show the 17% protection target: when the right line falls within light blue, protection of the study region is $< 17\%$. Gray regions in the map correspond to other forest hotspots considered for analysis but with less than 1000 eBird checklists (after data selection). The black lines in the map indicate the limits of each continent used to calculate observer experience and expertise.

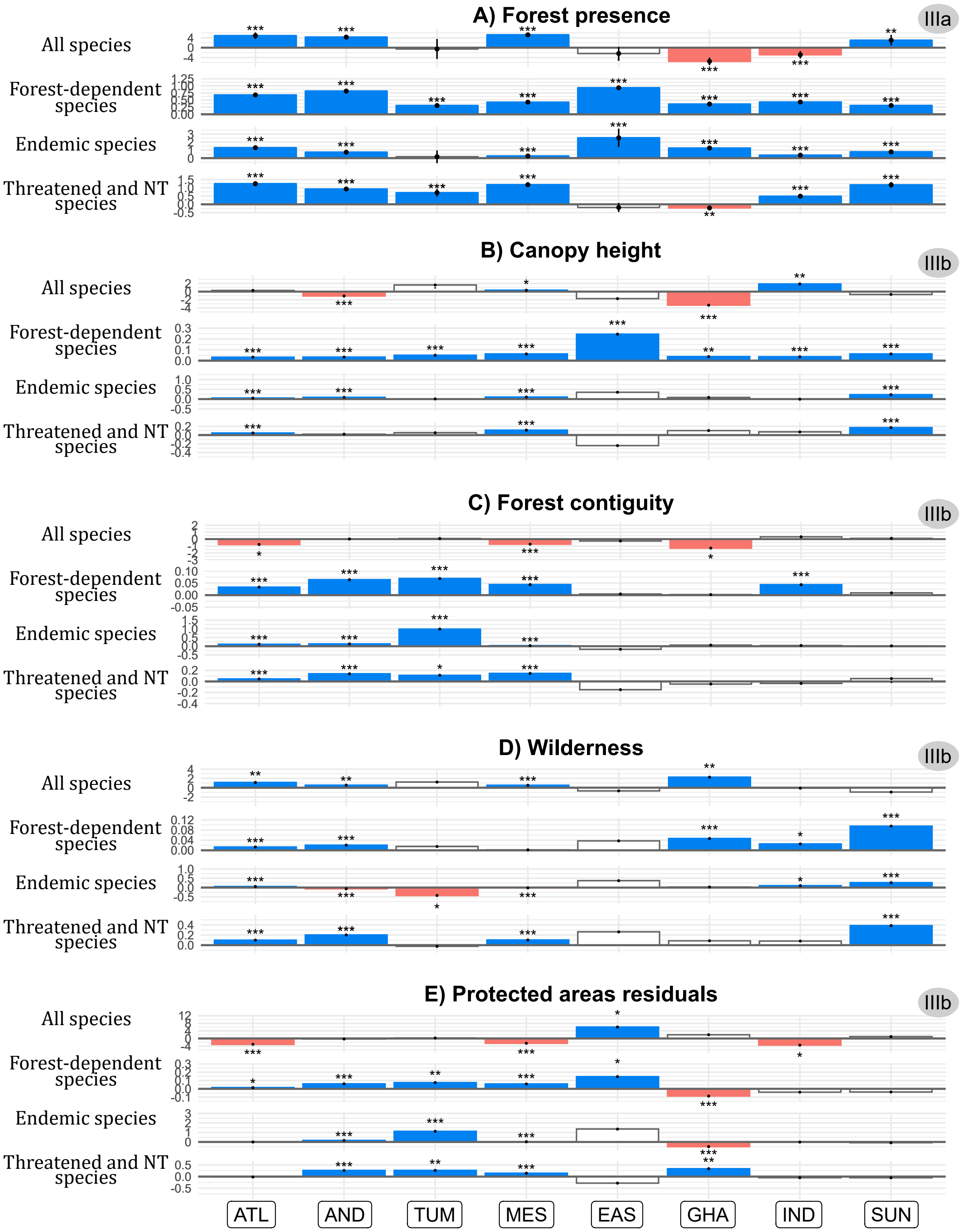


Extended Figure 2: Effects of forest presence and forest quality on bird diversity per hotspot. Analysis IIIa (A) shows the effect of forest presence on bird diversity. Analysis IIIb shows the effect of forest quality (B, Canopy height; C, Forest contiguity; D, Wilderness; E, Protected areas residuals) on bird diversity. Bird diversity is measured through four indices of richness in: all species, forest-dependent species, endemic species, threatened and Near Threatened species. Coefficients correspond to the estimates of GAM models; significance given by P-value $(* * * < 0.001 < ** < 0.10 < * < 0.05)$, and 95% confidence interval (vertical lines). Hotspots: ATL (Atlantic Forest), AND (Tropical Andes), TUM (Tumbes-Chocó-Magdalena), MES (Mesoamerica), EAS (Eastern Afromontane), GHA (Western Ghats and Sri Lanka), IND (Indo-Burma), SUN (Sundaland).

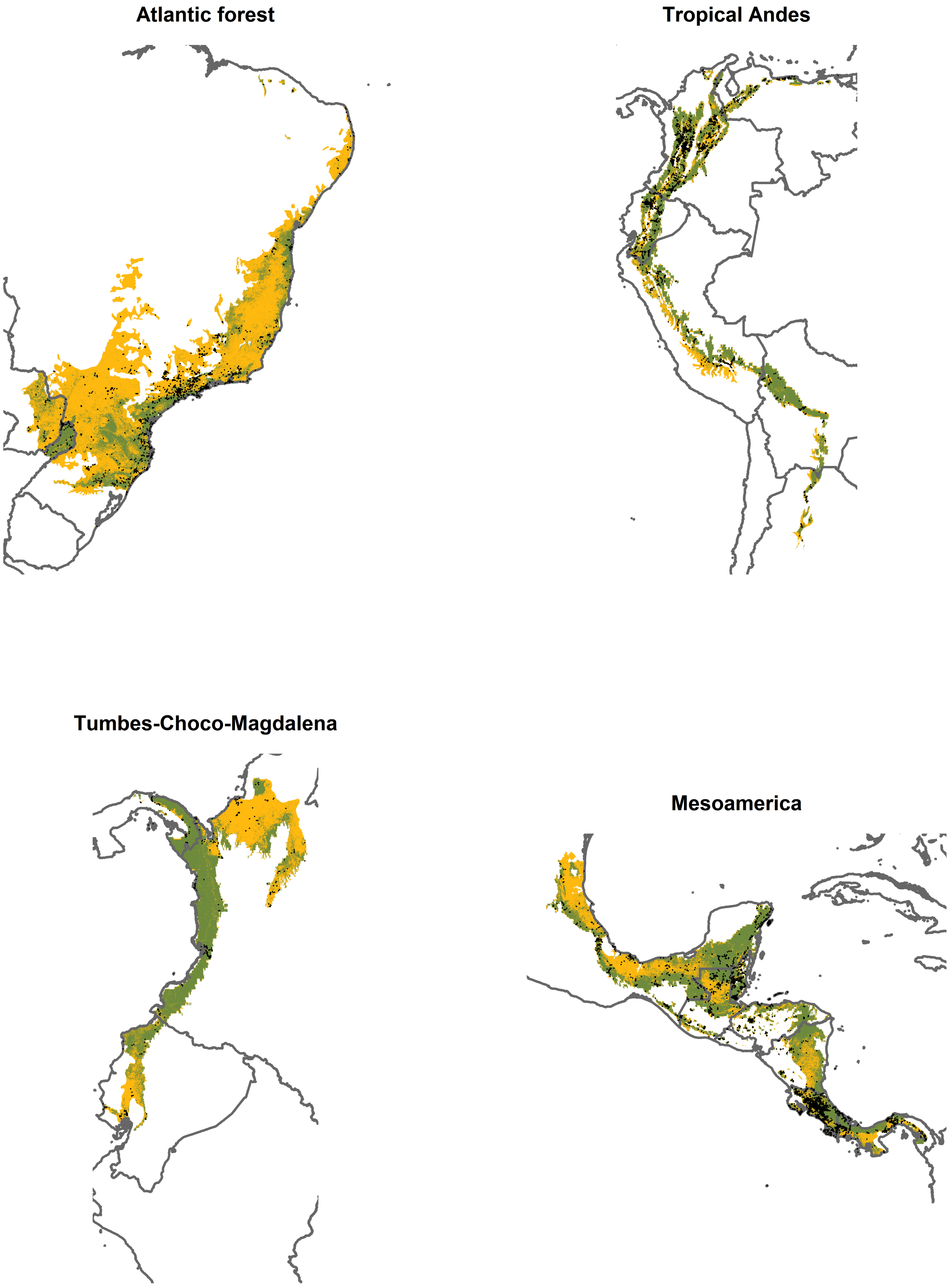


Extended Figure 3: Extent and sampling effort within each of the hotspot in the Americas. Each map indicates the full extent of the area analysed (i.e., the intersection between the hotspot and the "tropical and subtropical moist broadleaf forests" biome). Green represents forest, orange other habitats. Black dots represent the locations of the checklists used in the analyses.

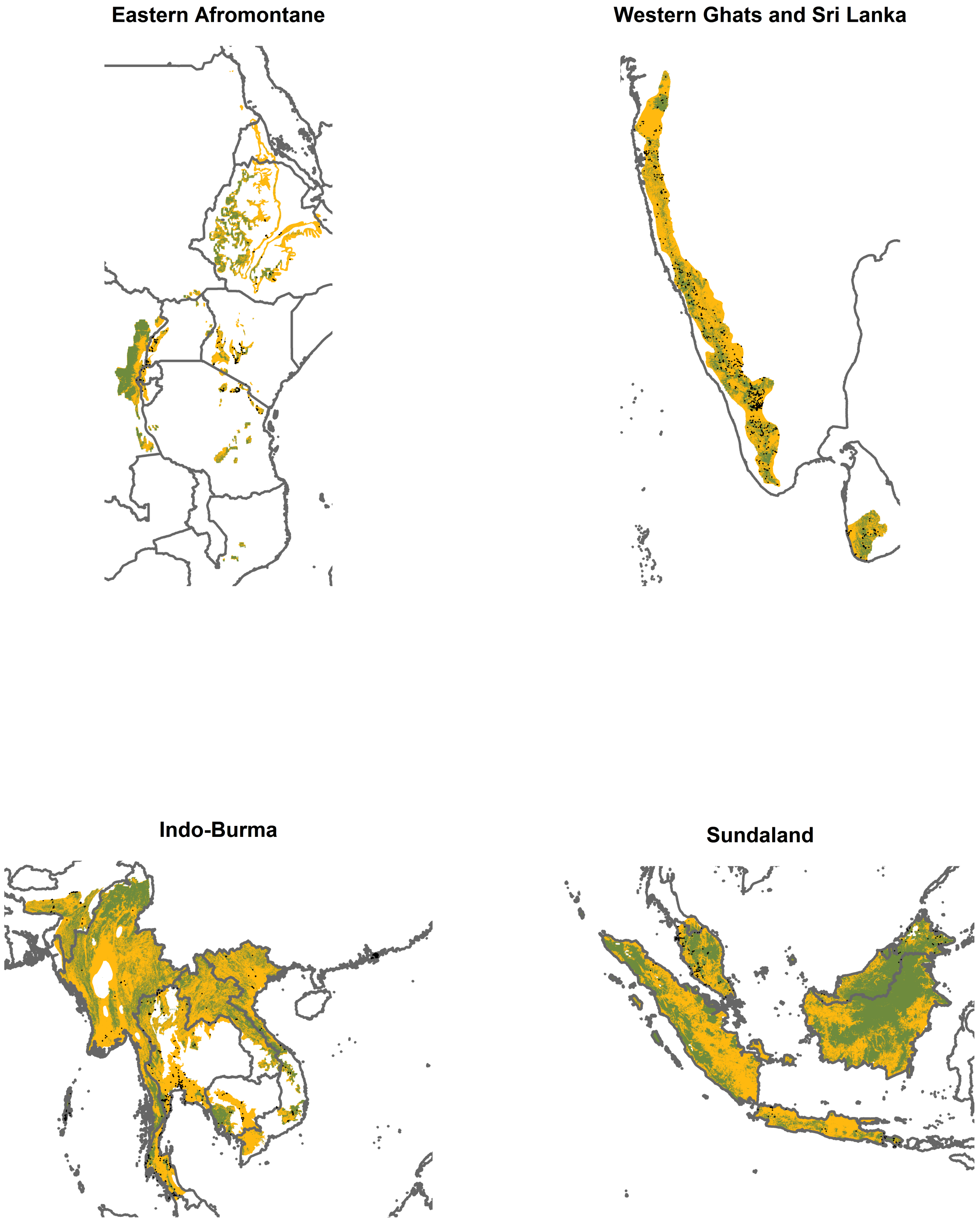


Extended Figure 4: Extent and sampling effort within each of the hotspot in Africa and Asia. Each map indicates the full extent of the area analysed (i.e., the intersection between the hotspot and the "tropical and subtropical moist broadleaf forests" biome). Green represents forest, orange other habitats. Black dots represent the locations of the checklists used in the analyses.

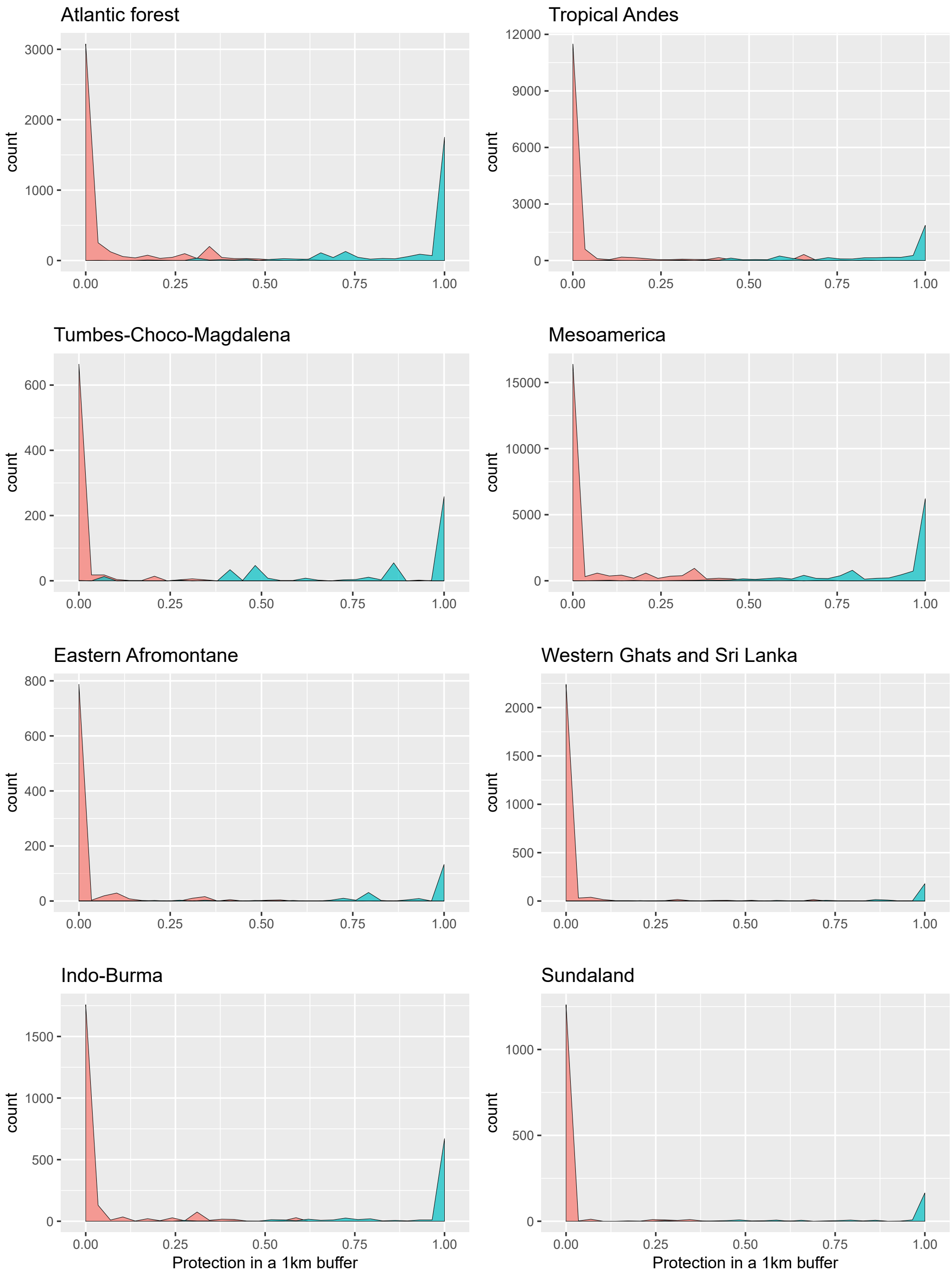


Extended Figure 5: Comparison between a binary classification of protection based on the protection status of the site coordinates (red for unprotected, blue for protected sites) as used in the analyses, and a continuous classification by measuring the proportion of area protected within a 1km buffer around each site.

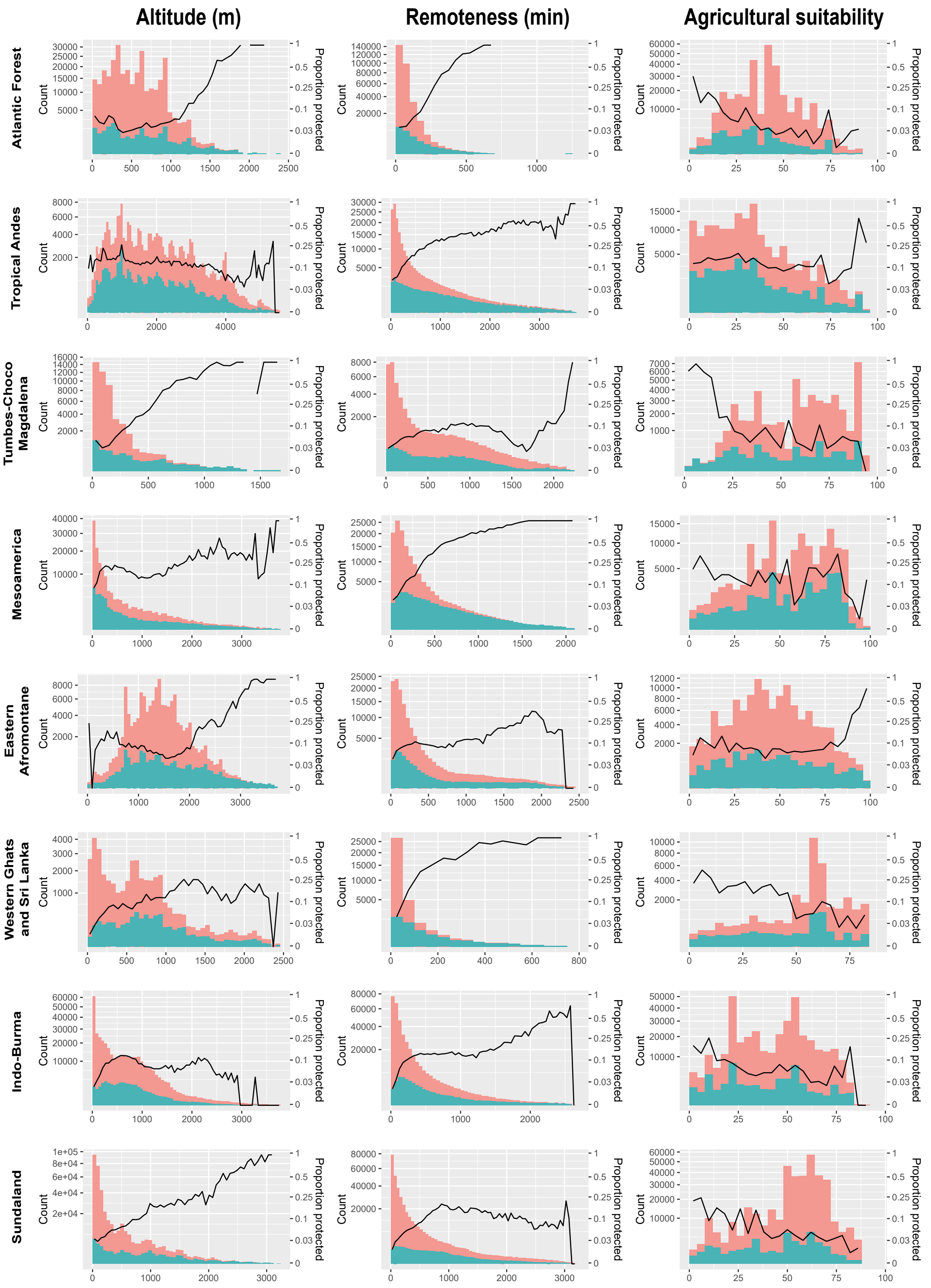


Extended Figure 6: Distribution of three variables used to control for location bias (altitude, remoteness, and agricultural suitability) for protected and unprotected sites (respectively blue and red bars) and proportion of sites protected for each bar (black lines), per hotspot. y-scales are transformed through square root function.

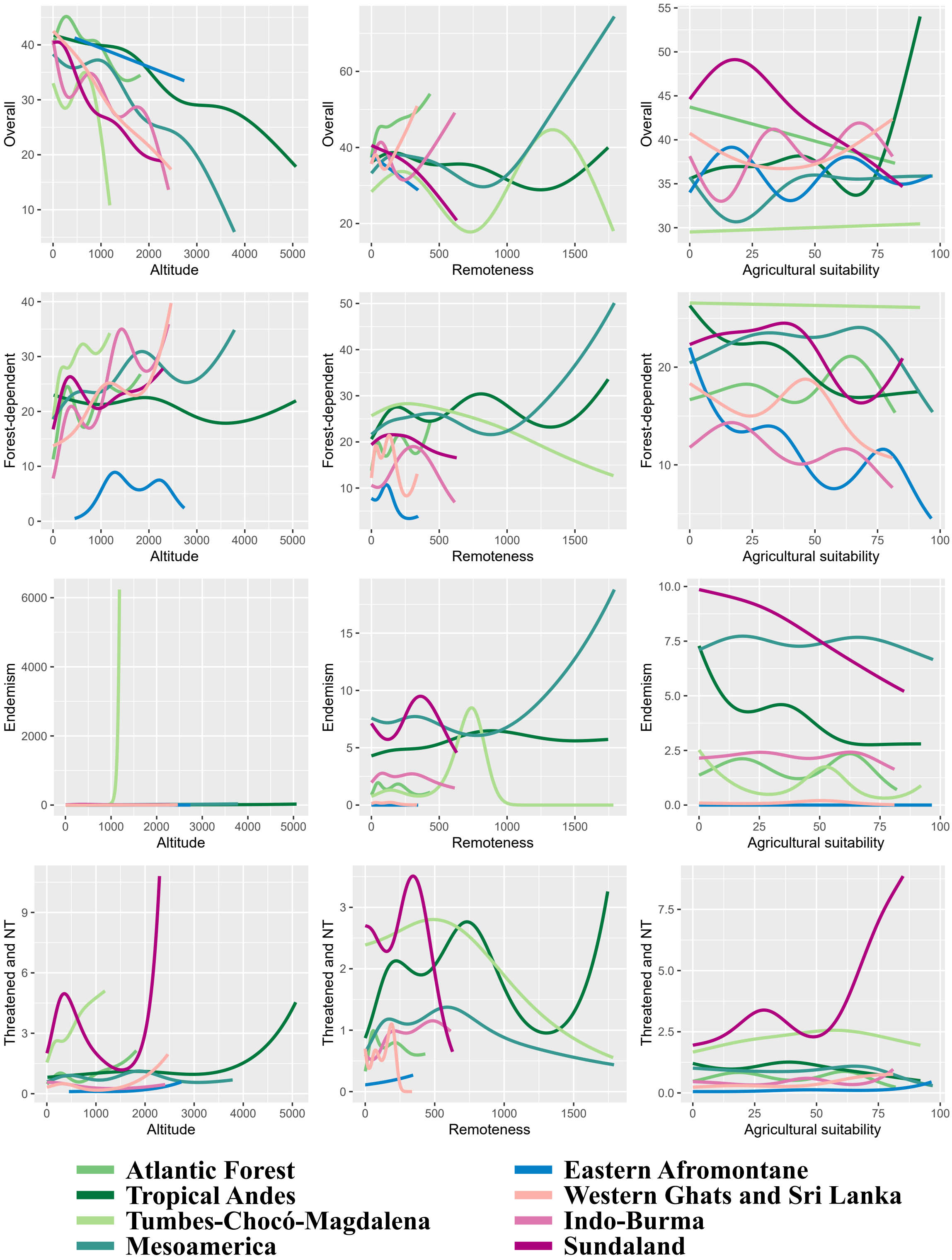


Extended Figure 7: Effects of each of the variables used to control for location bias in analysis I (altitude, remoteness and agricultural suitability) on each of the indices of bird diversity (overall species richness, richness in forest-dependent species, richness in endemic species, and richness in threatened and Near Threatened species), for each hotspot. Effects were predicted fixing all other variables to their median values and the protection variable to "unprotected".

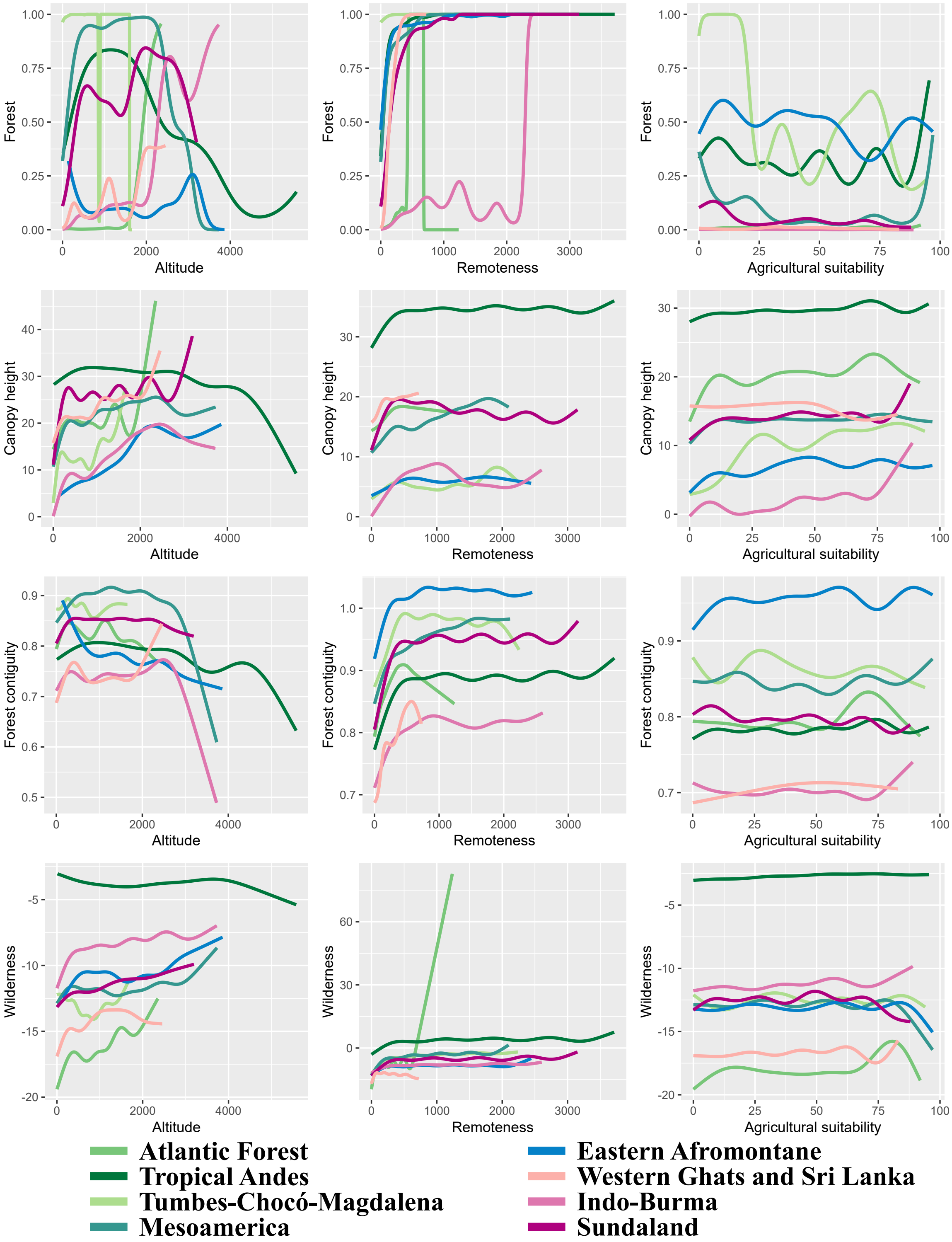


Extended Figure 8: Effects of each of the variables used to control for location bias in analysis II (altitude, remoteness and agricultural suitability) on forest presence and on each of the indices of forest quality (canopy height, forest contiguity, wilderness), for each hotspot. Effects were predicted fixing all other variables to their median values and the protection variable to "unprotected".

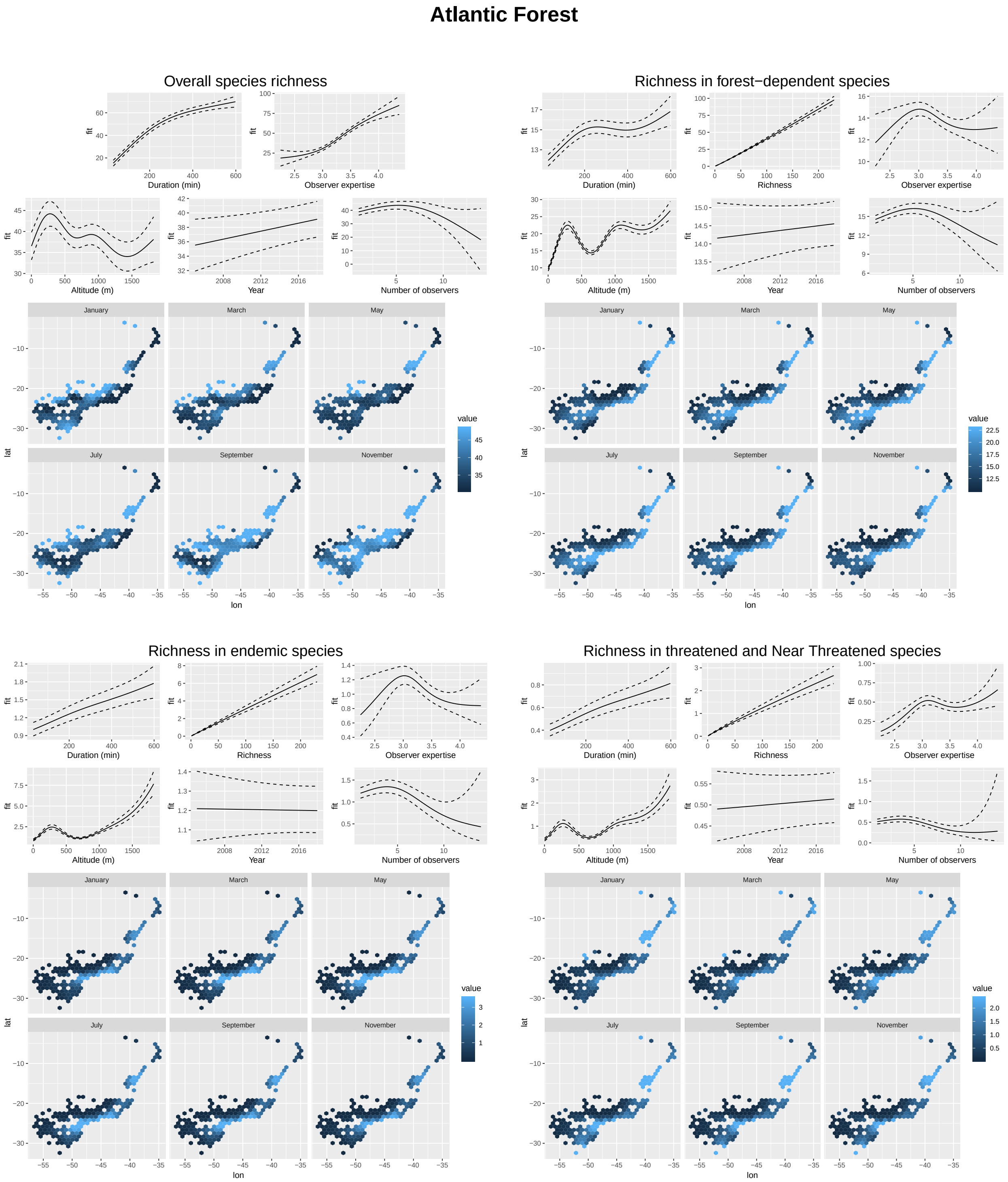


Extended Figure 9: Effects of each of the covariates used as controls in analysis III on each of the four bird diversity indices (overall species richness, richness in forest-dependent species, richness in endemic species, and richness in threatened and Near Threatened species), for the Atlantic Forest hotspot. We predicted bird indices (i.e., y values are always number of species) fixing all other variables to their median values. Maps represent spatial variation and seasonality for each diversity index. They correspond to a predict of each bird diversity index obtained by making longitude and latitude vary across the hotspot and fixing other variables to their median values, for 6 dates (mid-January [day 15], mid-March [day 74], mid-May [day 135], mid-July [day 196], mid-September [day 258], mid-November [day 319]), smoothed on a hexagonal grid by the ggplot function *stat_summary_hex* with default settings.

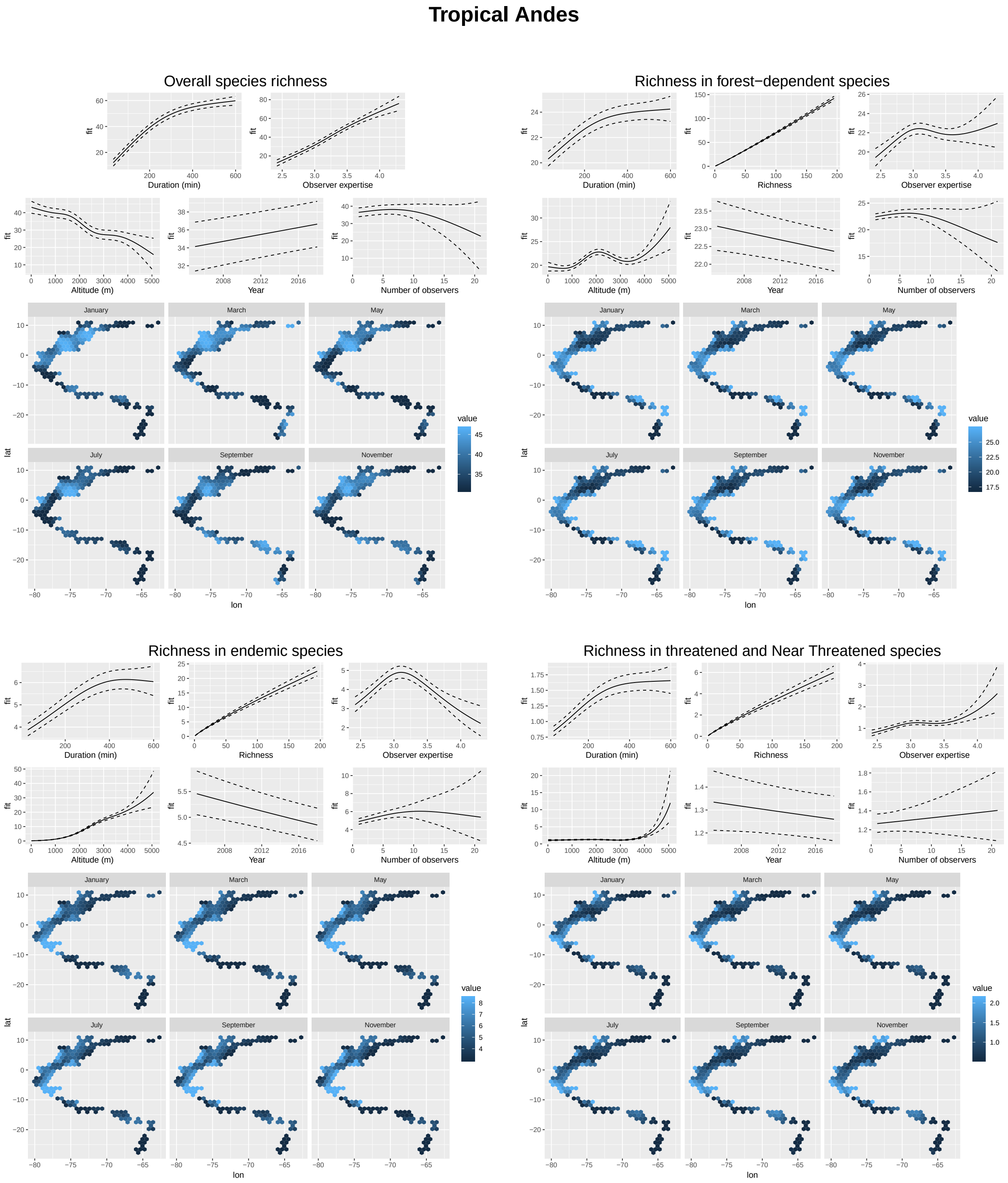


Extended Figure 10: Effects of each of the covariates used as controls in analysis III on each of the four bird diversity indices (overall species richness, richness in forest-dependent species, richness in endemic species, and richness in threatened and Near Threatened species), for the Tropical Andes hotspot. We predicted bird indices (i.e., y values are always number of species) fixing all other variables to their median values. Maps represent spatial variation and seasonality for each diversity index. They correspond to a predict of each bird diversity index obtained by making longitude and latitude vary across the hotspot and fixing other variables to their median values, for 6 dates (mid-January [day 15], mid-March [day 74], mid-May [day 135], mid-July [day 196], mid-September [day 258], mid-November [day 319]), smoothed on a hexagonal grid by the ggplot function *stat_summary_hex* with default settings.

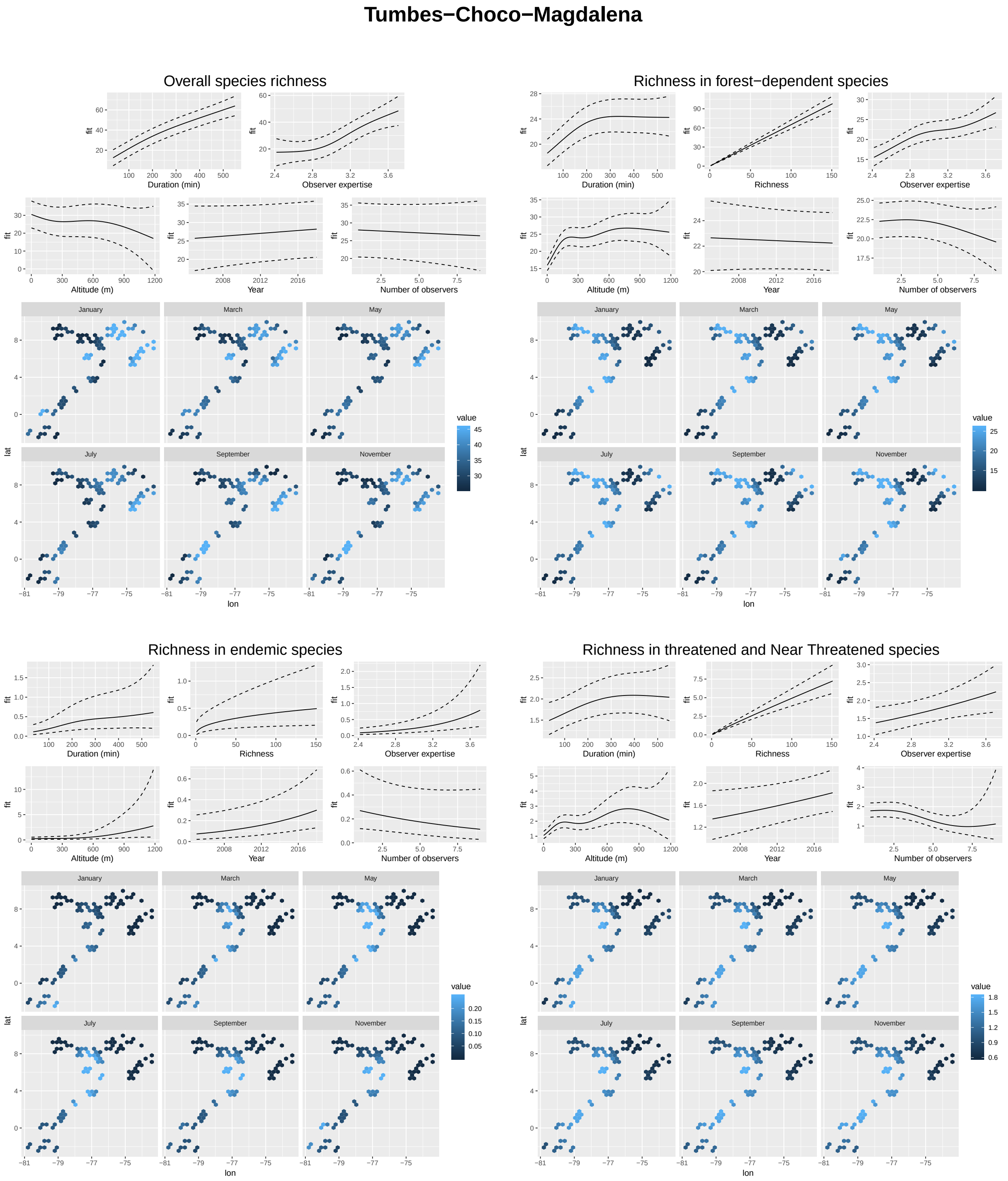


Extended Figure 11: Effects of each of the covariates used as controls in analysis III on each of the four bird diversity indices (overall species richness, richness in forest-dependent species, richness in endemic species, and richness in threatened and Near Threatened species), for the Tumbes-Choco-Magdalena hotspot. We predicted bird indices (i.e., y values are always number of species) fixing all other variables to their median values. Maps represent spatial variation and seasonality for each diversity index. They correspond to a predict of each bird diversity index obtained by making longitude and latitude vary across the hotspot and fixing other variables to their median values, for 6 dates (mid-January [day 15], mid-March [day 74], mid-May [day 135], mid-July [day 196], mid-September [day 258], mid-November [day 319]), smoothed on a hexagonal grid by the ggplot function *stat_summary_hex* with default settings.

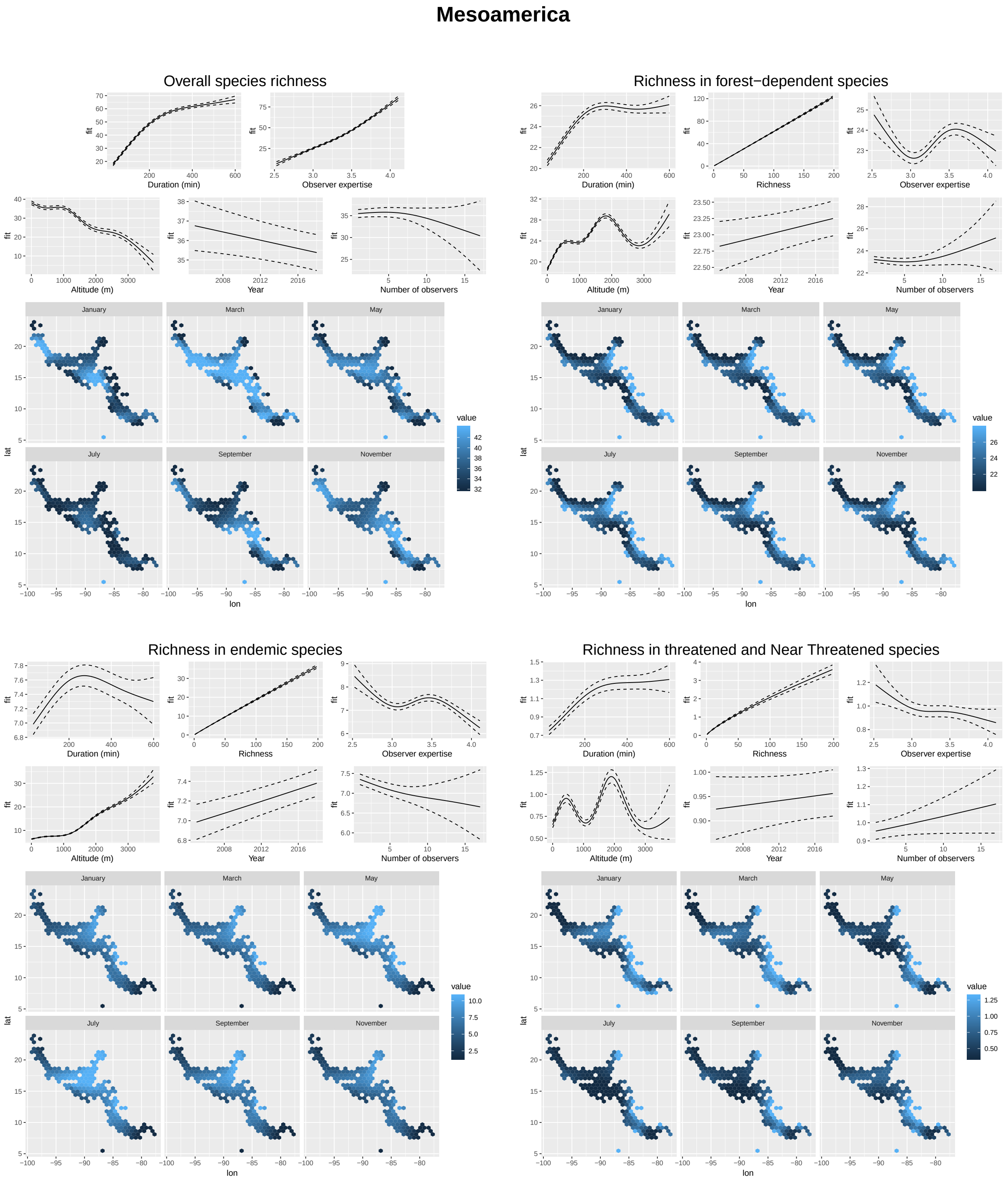


Extended Figure 12: Effects of each of the covariates used as controls in analysis III on each of the four bird diversity indices (overall species richness, richness in forest-dependent species, richness in endemic species, and richness in threatened and Near Threatened species), for the Mesoamerica hotspot. We predicted bird indices (i.e., y values are always number of species) fixing all other variables to their median values. Maps represent spatial variation and seasonality for each diversity index. They correspond to a predict of each bird diversity index obtained by making longitude and latitude vary across the hotspot and fixing other variables to their median values, for 6 dates (mid-January [day 15], mid-March [day 74], mid-May [day 135], mid-July [day 196], mid-September [day 258], mid-November [day 319]), smoothed on a hexagonal grid by the ggplot function *stat_summary_hex* with default settings.

### Eastern Afromontane

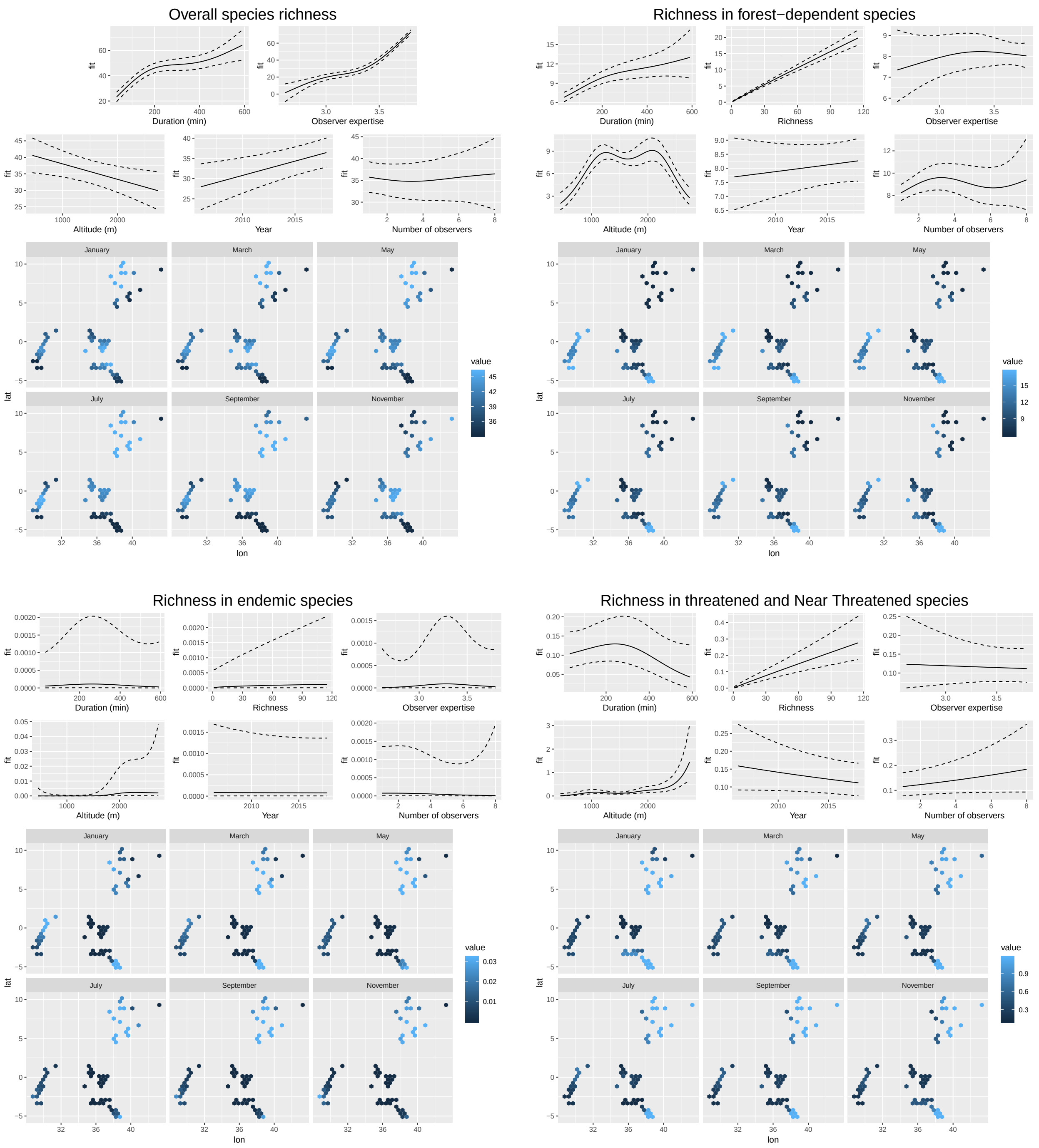


Extended Figure 13: Effects of each of the covariates used as controls in analysis III on each of the four bird diversity indices (overall species richness, richness in forest-dependent species, richness in endemic species, and richness in threatened and Near Threatened species), for the Eastern Afromontane hotspot. We predicted bird indices (i.e., y values are always number of species) fixing all other variables to their median values. Maps represent spatial variation and seasonality for each diversity index. They correspond to a predict of each bird diversity index obtained by making longitude and latitude vary across the hotspot and fixing other variables to their median values, for 6 dates (mid-January [day 15], mid-March [day 74], mid-May [day 135], mid-July [day 196], mid-September [day 258], mid-November [day 319]), smoothed on a hexagonal grid by the ggplot function *stat_summary_hex* with default settings.

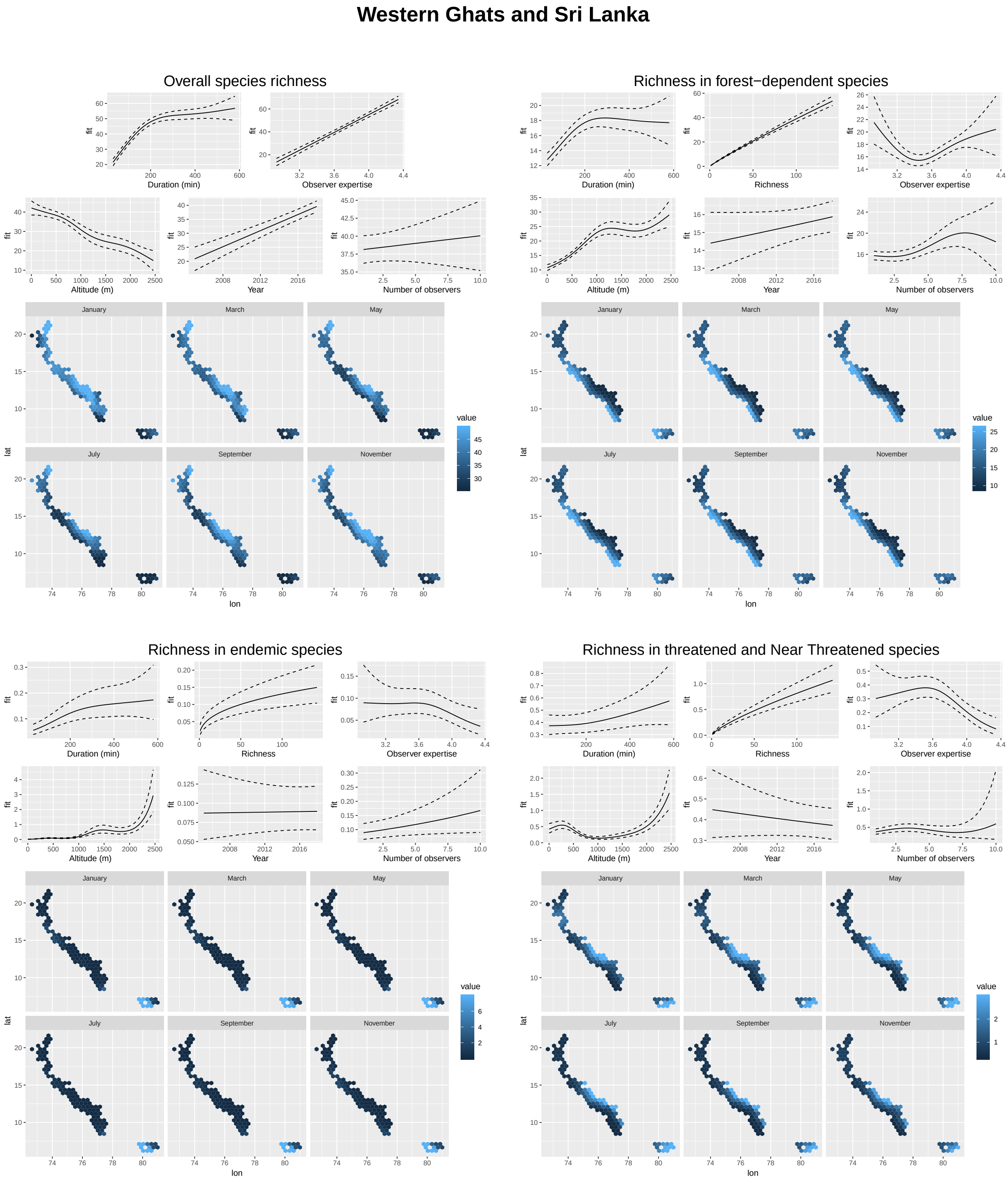


Extended Figure 14: Effects of each of the covariates used as controls in analysis III on each of the four bird diversity indices (overall species richness, richness in forest-dependent species, richness in endemic species, and richness in threatened and Near Threatened species), for the Western Ghats and Sri Lanka hotspot. We predicted bird indices (i.e., y values are always number of species) fixing all other variables to their median values. Maps represent spatial variation and seasonality for each diversity index. They correspond to a predict of each bird diversity index obtained by making longitude and latitude vary across the hotspot and fixing other variables to their median values, for 6 dates (mid-January [day 15], mid-March [day 74], mid-May [day 135], mid-July [day 196], mid-September [day 258], mid-November [day 319]), smoothed on a hexagonal grid by the ggplot function *stat_summary_hex* with default settings.

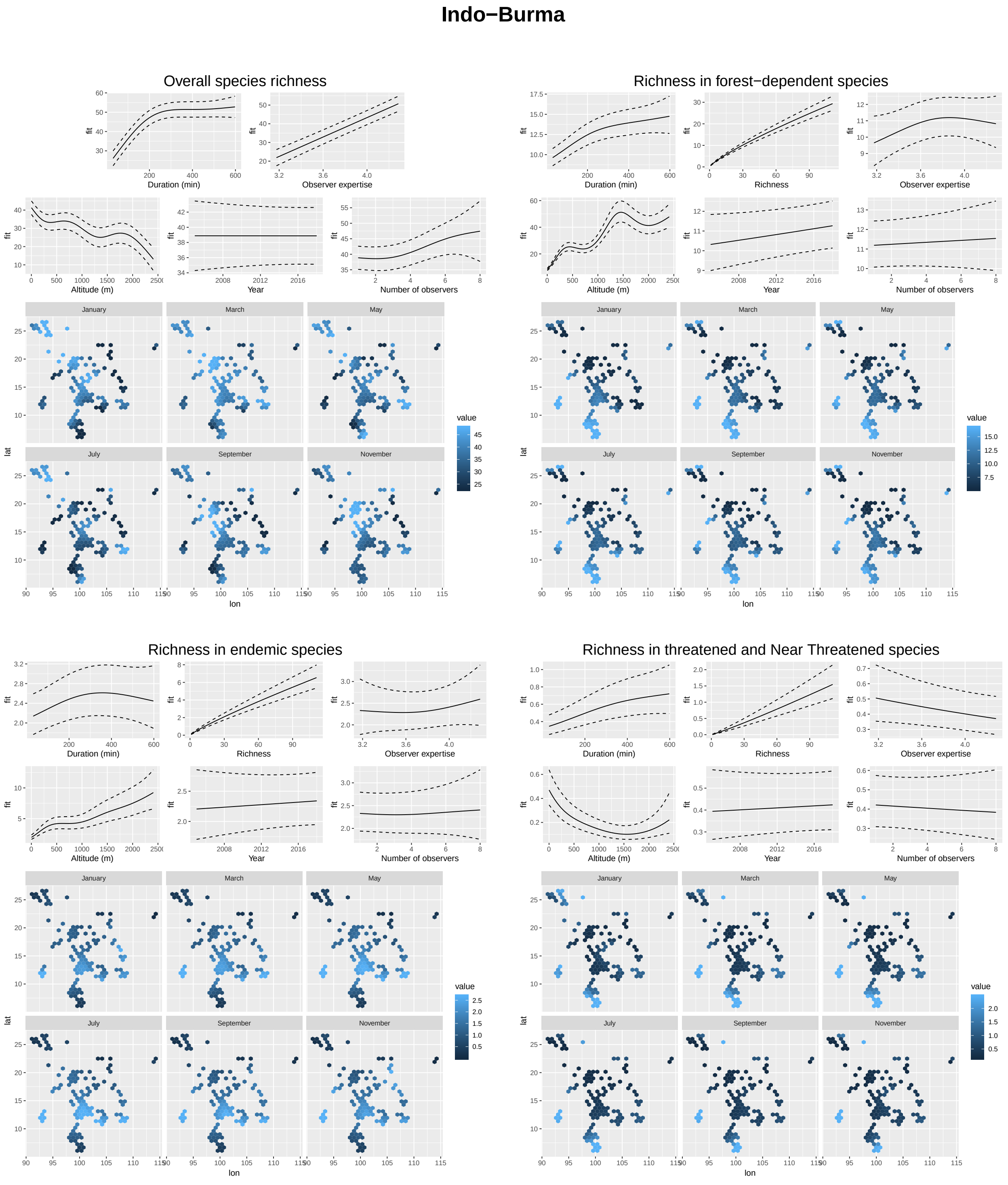


Extended Figure 15: Effects of each of the covariates used as controls in analysis III on each of the four bird diversity indices (overall species richness, richness in forest-dependent species, richness in endemic species, and richness in threatened and Near Threatened species), for the Indo-Burma hotspot. We predicted bird indices (i.e., y values are always number of species) fixing all other variables to their median values. Maps represent spatial variation and seasonality for each diversity index. They correspond to a predict of each bird diversity index obtained by making longitude and latitude vary across the hotspot and fixing other variables to their median values, for 6 dates (mid-January [day 15], mid-March [day 74], mid-May [day 135], mid-July [day 196], mid-September [day 258], mid-November [day 319]), smoothed on a hexagonal grid by the ggplot function *stat_summary_hex* with default settings.

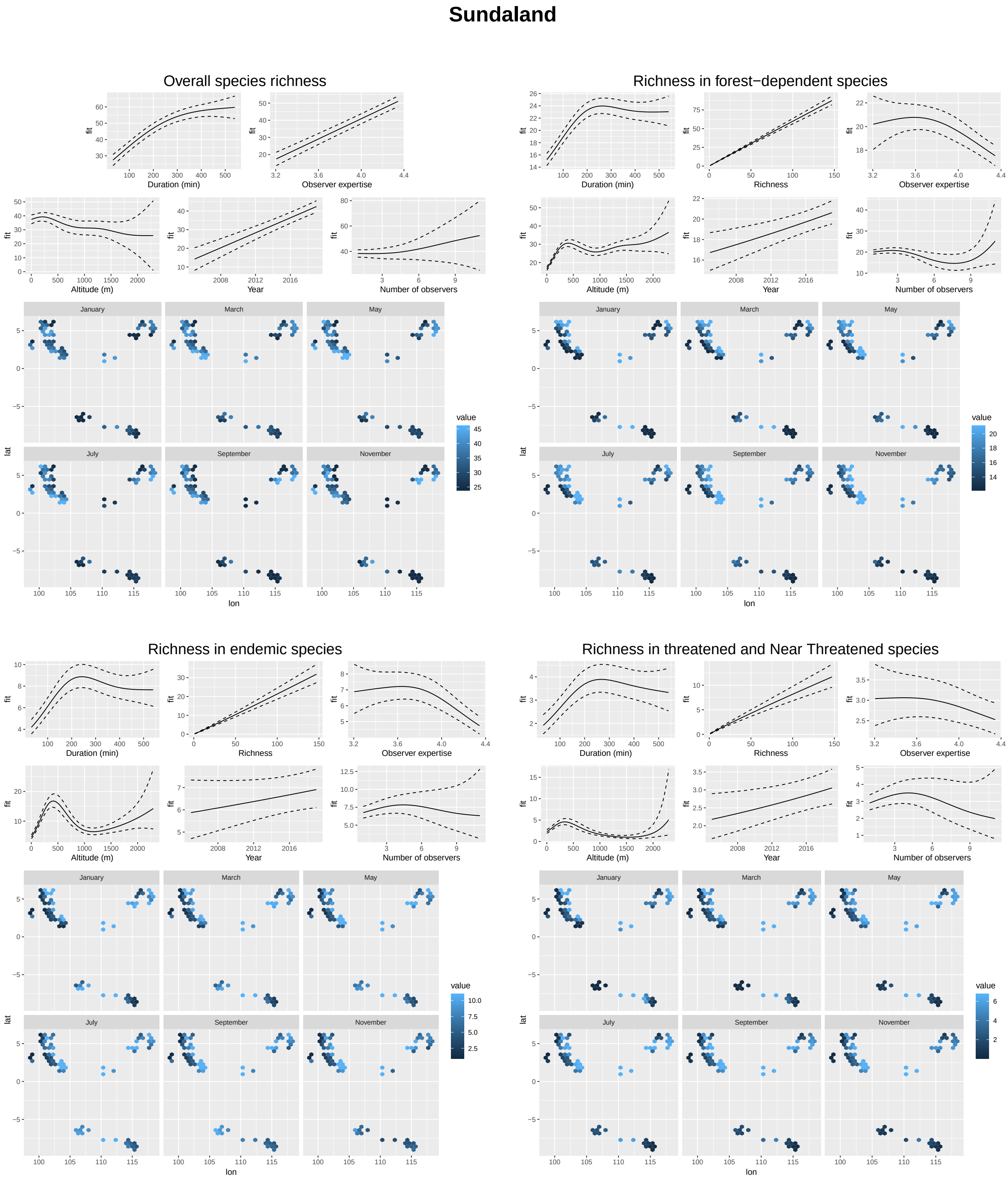


Extended Figure 16: Effects of each of the covariates used as controls in analysis III on each of the four bird diversity indices (overall species richness, richness in forest-dependent species, richness in endemic species, and richness in threatened and Near Threatened species), for the Sundaland hotspot. We predicted bird indices (i.e., y values are always number of species) fixing all other variables to their median values. Maps represent spatial variation and seasonality for each diversity index. They correspond to a predict of each bird diversity index obtained by making longitude and latitude vary across the hotspot and fixing other variables to their median values, for 6 dates (mid-January [day 15], mid-March [day 74], mid-May [day 135], mid-July [day 196], mid-September [day 258], mid-November [day 319]), smoothed on a hexagonal grid by the ggplot function *stat_summary_hex* with default settings.
