## Extended tables for "Effectiveness of protected areas in conserving tropical forest birds"

**Extended Table 1:** Effect of protected areas on each of the response variables considered in analyses I, II and IIIa, measured as percentage of difference. For analysis I, this was obtained by first predicting the response variable in a protected [Resp_In_] and in an unprotected site [Resp_Out_], while fixing all other variables to their median value. We then calculated the percentage of increase due to protection [*100 * (*Resp_In_ *–* Resp_Out_*) / abs(*Resp_Out_*)*], which estimates how richer an average site can be if protected rather than unprotected. We did the same for analysis IIIa, predicting the response variables in two unprotected sites, one forested [Resp_In_] and one not forested [Resp_Out_], with all variables fixed to their median values. For analysis II, we focused on background sites that are currently protected. We then predicted from our models response variables (probability of forest presence; each of the three habitat quality variables) for each site, first setting them as protected [Resp_PA_], and second setting them as unprotected [Resp_unPA_]. We then calculated the ratio percentage of increase due to protection [*100 * (*Resp_PA_ *–* Resp_unPA_*) / abs(*Resp_unPA_*)*] which estimates how much habitat loss or degradation would have happened had these sites not been protected. Column “MEAN” shows the average effect across the eight hotspots.

| **AnalysIS** | **Response variable** | ATL  (%) | AND  (%) | TUM  (%) | MES  (%) | EAS  (%) | GHA  (%) | IND  (%) | SUN  (%) | **Mean**  (%) |
| --- | --- | --- | --- | --- | --- | --- | --- | --- | --- | --- |
| **I** | Overall Richness | – 3.2 | 0.0 | 11.4 | – 5.1 | 2.4 | – 1.3 | 2.0 | 3.4 | **1.7** |
| **I** | Forest-dependent | 22.8 | 13.4 | 5.1 | 13.8 | 78.7 | 6.8 | 1.5 | 0.1 | **17.8** |
| **I** | Endemic | 18.9 | 24.0 | – 24.2 | 7.1 | 635.4 | – 25.5 | – 6.4 | – 8.6 | **77.6** |
| **I** | Threatened and Near Threatened | 17.1 | 33.7 | 7.3 | 37.1 | – 22.0 | 27.7 | 58.9 | – 7.5 | **19.0** |
| **II** | Forest presence | 51.6 | 3.6 | 1.7 | 4.2 | 32.9 | 18.1 | 20.2 | 10.1 | **17.8** |
| **II** | Canopy height | 5.1 | -0.5 | 15.0 | 3.1 | 3.8 | -1.5 | 10.5 | 2.8 | **4.8** |
| **II** | Forest contiguity | 5.4 | 0.9 | 1.4 | 2.2 | 0.8 | 3.5 | 4.6 | 1.7 | **2.6** |
| **II** | Wilderness | 1.3 | 2.2 | 6.4 | 4.5 | 3.0 | 4.0 | 9.0 | 14.9 | **5.7** |
| **IIIa** | Overall Richness | 13.0 | 13.1 | -1.8 | 17.3 | -6.5 | -14.2 | -7.2 | 8.2 | **2.7** |
| **IIIa** | Forest-dependent | 97.3 | 126.0 | 35.9 | 53.0 | 153.8 | 43.2 | 53.8 | 36.1 | **74.9** |
| **IIIa** | Endemic | 266.9 | 109.2 | 18.2 | 30.5 | 1160.1 | 252.7 | 44.4 | 118.0 | **250.0** |
| **IIIa** | Threatened and Near Threatened | 246.7 | 152.0 | 101.8 | 227.1 | -17.3 | -19.6 | 63.0 | 223.1 | **122.1** |

**Extended Table 2:** Summary statistics per hotspot. Number of eBird checklists, observations, species and observers, after data selection (i.e. as used in the analyses). Average values for each of the four bird diversity indices considered: overall richness, richness in forest-dependent species, richness in endemics, and richness in threatened and Near Threatened species.

| Hotspot name | Hotspot code | Number of checklists | Number of observations | Number of species | Number of observers | Richness  (median ± se) | Richness in forest-dependent species  (median ± se) | Richness in endemic species  (median ± se) | Richness in threatened and Near Threatened species  (median ± se) |
| --- | --- | --- | --- | --- | --- | --- | --- | --- | --- |
| **Atlantic Forest** | ATL | 6,760 | 286,547 | 940 | 928 | 38 ± 26 | 19 ± 21 | 1 ± 4 | 1 ± 3 |
| **Tropical Andes** | AND | 17,758 | 683,213 | 2,229 | 2,244 | 34 ± 24 | 22 ± 19 | 4 ± 9 | 1 ± 2 |
| **Tumbes-Chocó-Magdalena** | TUM | 1,188 | 44,382 | 914 | 509 | 34 ± 21 | 16 ± 17 | 0 ± 0 | 1 ± 1 |
| **Mesoamerica** | MES | 32,784 | 1,363,889 | 1,185 | 3503 | 38 ± 24 | 24 ± 18 | 6 ± 6 | 1 ± 1 |
| **Eastern Afromontane** | EAS | 1,097 | 52,364 | 986 | 263 | 46 ± 25 | 11 ± 9 | 0 ± 0 | 0 ± 1 |
| **Western Ghats and Sri Lanka** | GHA | 2,646 | 99,567 | 487 | 556 | 37 ± 19 | 15 ± 11 | 0 ± 2 | 0 ± 1 |
| **Indo-Burma** | IND | 2,996 | 102,558 | 1,031 | 418 | 34 ± 17 | 10 ± 12 | 1 ± 2 | 0 ± 2 |
| **Sundaland** | SUN | 1,548 | 54,534 | 706 | 170 | 33 ± 19 | 15 ± 16 | 2 ± 11 | 1 ± 5 |
