## Supplementary discussion for "Effectiveness of protected areas in conserving tropical forest birds"

When comparing across hotspots, we found some heterogeneity in our results, potentially induced by three factors. First, failure to comply with our simplifying hypothesis that our study regions (i.e. extent of biodiversity hotspots included in the “tropical and subtropical moist broadleaf forests” biome) were originally covered by homogeneous forest could explain some differences in bird responses. For instance, responses of bird diversity indices in Western Ghats and Sri Lanka were often low, sometimes the opposite of others. This could be due to the natural habitat heterogeneity of this region, such as the presence of natural grasslands above the shola forests and large variations in rainfall patterns (Mittermeier, 2004); this may also be the case in the Tropical Andes, which include high natural grasslands such as Páramo (see Supplementary Methods 4D for further information on this assumption), and perhaps Eastern Afromontane. This corroborates with the high proportion of species with null dependency of forest in Western Ghats and Sri Lanka, as well as Eastern Afromontane (respectively 33% and 38% of species detected, against an average of 18% for other hotspots). Second, differences in protection regimes could explain some of the variation found between hotspots. Indeed, the location of protected areas in the Atlantic Forest is not highly biased towards remote and high areas (reactive approach (Brooks et al., 2006), Extended Figure 6), which could explain the high effectiveness measured. Conversely, protected areas in Sundaland are highly biased towards remote and high zones that are less likely to suffer from human pressure in the short-term (pro-active approach (Brooks et al., 2006), Extended Figure 6), which could explain the low effect we measured while controlling for location biases. Finally, sampling effort greatly differed between hotspots (ranging from 1,070 checklists analysed in Eastern Afromontane to 31,053 in Mesoamerica), which could affect the statistical power of tests. This could explain why hotspots in the Americas showed clearer results than those from Asia and Africa, particularly for the effects of protected areas in maintaining forest quality. The consistency of results we got across continents in analyses II and III (disentangling the mechanisms of protected area effects on bird diversity) give high credit to this assumption.

**References**

Brooks, T.M., Mittermeier, R.A., da Fonseca, G.A.B., Gerlach, J., Hoffmann, M., Lamoreux, J.F., Mittermeier, C.G., Pilgrim, J.D., and Rodrigues, A.S.L. (2006). Global Biodiversity Conservation Priorities. Science *313*, 58–61.

Mittermeier, R.A. (2004). Hotspot revisited (Cemex).
